# Dimensionality in recurrent spiking networks: global trends in activity and local origins in connectivity

**DOI:** 10.1101/394684

**Authors:** Stefano Recanatesi, Gabriel Koch Ocker, Michael A. Buice, Eric Shea-Brown

## Abstract

The dimensionality of a network’s collective activity is of increasing interest in neuroscience. This is because dimensionality provides a compact measure of how coordinated network-wide activity is, in terms of the number of modes (or degrees of freedom) that it can independently explore. A low number of modes suggests a compressed low dimensional neural code and reveals interpretable dynamics [1], while findings of high dimension may suggest flexible computations [2, 3]. Here, we address the fundamental question of how dimensionality is related to connectivity, in both autonomous and stimulus-driven networks. Working with a simple spiking network model, we derive three main findings. First, the dimensionality of global activity patterns can be strongly, and systematically, regulated by local connectivity structures. Second, the dimensionality is a better indicator than average correlations in determining how constrained neural activity is. Third, stimulus evoked neural activity interacts systematically with neural connectivity patterns, leading to network responses of either greater or lesser dimensionality than the stimulus.

**Author summary:** New recording technologies are producing an amazing explosion of data on neural activity. These data reveal the simultaneous activity of hundreds or even thousands of neurons. In principle, the activity of these neurons could explore a vast space of possible patterns. This is what is meant by high-dimensional activity: the number of degrees of freedom (or “modes”) of multineuron activity is large, perhaps as large as the number of neurons themselves. In practice, estimates of dimensionality differ strongly from case to case, and do so in interesting ways across experiments, species, and brain areas. The outcome is important for much more than just accurately describing neural activity: findings of low dimension have been proposed to allow data compression, denoising, and easily readable neural codes, while findings of high dimension have been proposed as signatures of powerful and general computations. So what is it about a neural circuit that leads to one case or the other? Here, we derive a set of principles that inform how the connectivity of a spiking neural network determines the dimensionality of the activity that it produces. These show that, in some cases, highly localized features of connectivity have strong control over a network’s global dimensionality—an interesting finding in the context of, e.g., learning rules that occur locally. We also show how dimension can be much different than first meets the eye with typical “pairwise” measurements, and how stimuli and intrinsic connectivity interact in shaping the overall dimension of a network’s response.

## 1 Introduction

A fundamental step toward understanding neural circuits is relating the structure of their dynamics to the structure of their connectivity [4, 5, 6, 7, 8, 9, 10, 11, 12, 13, 14]. However, the underlying networks are typically so complex that it is apriori unclear what features of the connectivity will matter most (and least) in driving network activity, and how the impacts of different connectivity features interact. Recent theoretical work has made progress in identifying rich and distinct roles for several different features of network connectivity: local connection structures [15, 16, 17, 18, 19, 20], spatial profiles of coupling [21], low-rank connection structures [22] and subnetwork statistics [23, 24].

Here we focus on linking network connectivity to collective activity as quantified by the dimensionality of the neural response. This dimensionality summarizes the number of collective modes, or degrees of freedom, that the network’s activity explores. We use the “participation ratio” dimension, which is directly computable from the pairwise covariances among all cells in a population [25, 3, 2, 26, 27]. This connection is useful because the structure of pairwise covariance has been linked, in turn, to the fidelity of the neural code, both at the single neuron [28], and at the population levels [29, 30, 31, 32, 33]. Overall, the participation ratio has proven useful in interpreting properties of multi-units neuronal recordings [26], and has yielded a remarkable perspective on neural plasticity and how high dimensional responses can be optimal for general computations [3, 2].

Two factors arise in our efforts to understand what it is about a network’s connectivity that determines the dimensionality of its activity. First, this process requires untangling two leading contributions to collective spiking: the reverberation of internal activity within the circuit, and its modulation by external inputs [34, 35, 36]. Experiments point out that both have strong effects [37, 38, 21, 39], and they interact in rich ways that our analysis will begin to dissect.

Second, beyond providing general formulas, the understanding we seek demands that we identify relatively simple “observables” of complex network connectivity that systematically determine the dimensionality they produce. A natural approach is based on connection paths through networks, and how these can in turn be decomposed into local circuit micro-circuits, or “motifs” [17, 18, 19, 20, 40, 41, 23, 16]. This is attractive because such local connectivity structures can be measured in tractable “multi-patch” type experiments, are limited in their complexity, and are controlled by local plasticity mechanisms. The prevalence of motifs, characterized in terms of connection probabilities and strengths, has achieved success in predicting the average levels of pairwise correlation among spiking cells – a measure of coordinated activity related to dimensionality in interesting ways that we will further explore below ([20, 17, 18]; see also [42]). Here we deploy this framework to compute the dimensionality of spontaneous and stimulus-driven neural activity. We find that expressions based on just the details of small (and hence local) connection motifs give correct qualitative, and in some (but not other) cases quantitative, predictions of trends in dimensionality of global activity patterns. This underlines the utility of local network motifs as building blocks in bridging from network connectomics to network dynamics.

Our main findings are threefold: First, the dimensionality of global activity patterns can be strongly, and systematically, regulated by local connectivity structures. Second, for a wide range of networks this dimensionality can be surprisingly low (indicating strongly coordinated activity) even when the average correlations among pairs of neurons are very weak, cfr.[43]. Third, the dimensionality of stimulus evoked neural activity is controlled systematically by neural connectivity, leading to network responses that have either expanded or reduced the dimension of the original stimulus.

In what follows we will start by introducing the underlying theoretical framework (Sec. 2.1). We describe the mathematical model, a spiking network of linearly interacting point process cells (a “Poisson linear network”, linearized GLM, or Hawkes process), together with the measure of dimensionality we use (Sec. 2.2). In Sec. 2.3, we show how this dimensionality can be expressed in terms of connectivity motifs. We continue by analyzing the dimensionality of the spontaneous (internally generated) activity of an excitatory randomly connected network (Sec. 2.4), and move to stimulus-driven networks of this type (Sec. 2.5). Finally, we generalize our results to consider different connectivity topologies as well as excitatory-inhibitory balanced networks (Sec. 2.6). We hinge the discussion around the question of how a network can modulate the dimensionality of its response to external stimuli by leveraging its local connectivity (Sec. 3).

## 2 Results

In recent years, neuroscientists have developed a flexible framework for predicting how spike train correlations are guided by the structure of recurrent connections [19, 17, 18, 42, 44]. Here we present and extend this framework to compute the dimensionality of spontaneous and stimulus-driven activity. The expert reader, who may be well acquainted with all this material, may be able to start reading from Sec. 2.3 or Sec. 2.4. In the same spirit we encourage the reader for whom the idea of network motifs is novel, to follow the more detailed presentation found in the Suppl. Mat. up to the result expressed in Eq. 11.

Throughout the paper bold lower-case letters will identify vectors, while bold upper case letters identify matrices. Non-bold letters identify scalar numbers.

### 2.1 The theoretical framework

Consider a recurrent neural network of *N* neurons where the activity *y*_*i*_(*t*) of neuron *i* at time *t* occurs around a baseline rate of irregular firing, which is set by the internal connectivity of the network ***W*** and an external input ***ξ***(*t*). The spike train of neuron *i* is given by 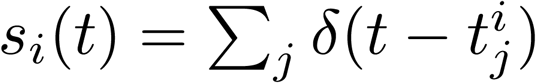 where each spike is sampled from a Poisson distribution with instantaneous mean rate (intensity) *y*_*i*_(*t*). The response of the whole network ***y***(*t*) can then be captured by linearizing its dynamics around the baseline rates, giving the equation:

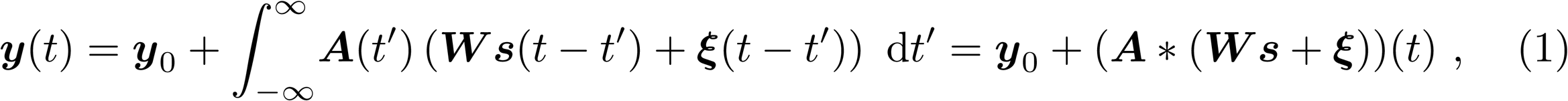

where each entry of the vector ***y***(*t*) is the instantaneous firing rate of neuron *i* at time *t* with baseline firing rate ***y***_0_. Here *W*_*ij*_ is the synaptic strength between neuron *i* and neuron *j*, and ***A*** is a diagonal matrix where *A*_*ii*_ is the postsynaptic filter which encapsulates the timecourse of the postsynaptic response. Thus, *G*_*ij*_ = *A*_*ii*_ *W*_*ij*_ defines an effective connectivity matrix. Finally, ***ξ*** is the external input to the network. This model is pictured in Fig. 1a, where the input ***ξ*** contributes to the baseline activity of each neuron, and the recurrent feedback is linearized.

**Fig 1.**
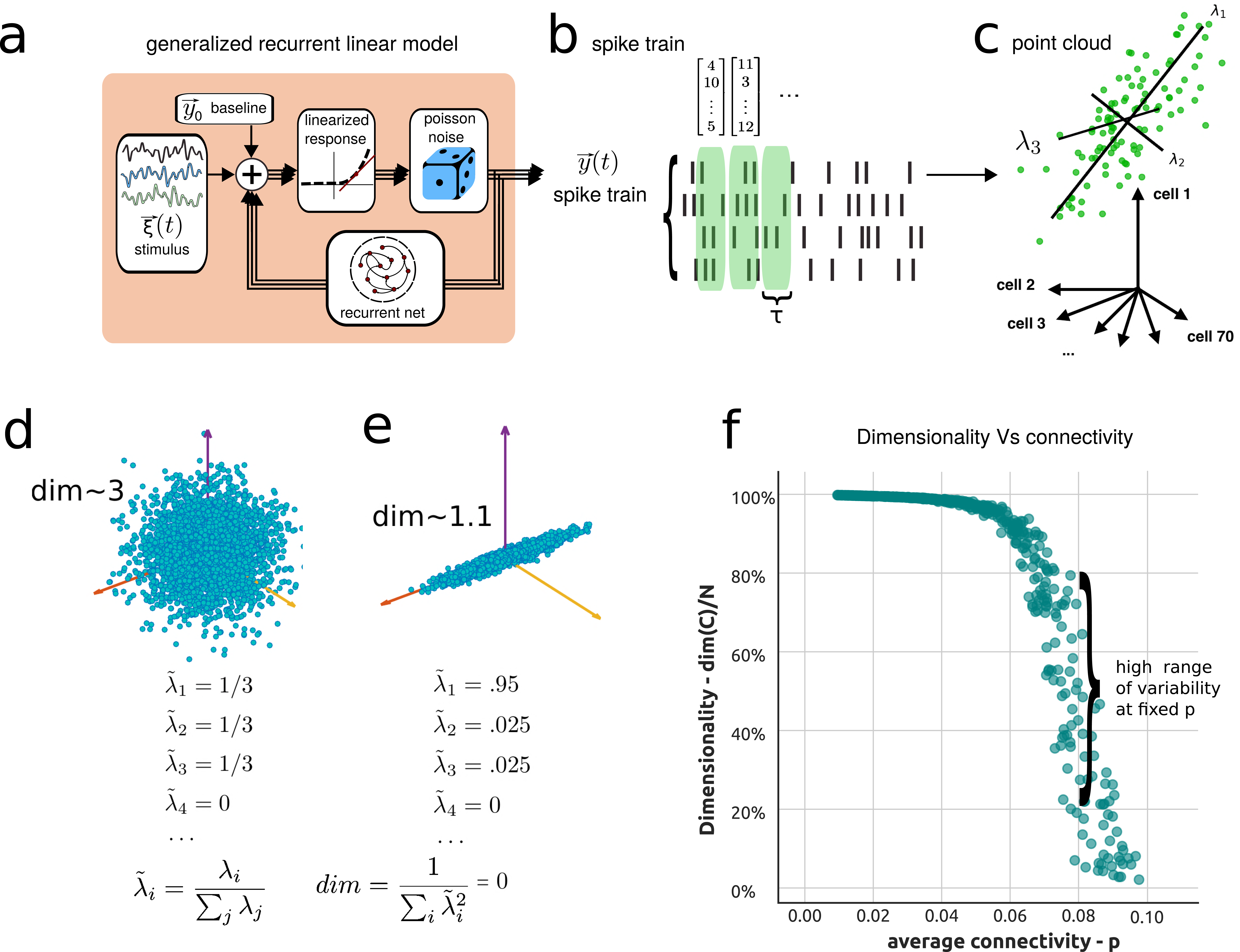
Dimensionality of the activity of a generalized linear recurrent network (a “linearized inhomogeneous Poisson GLM.” [45, 46]) **a**) Schematic of a generalized linear recurrent neural network. **b**) Spike train generated by the model, showing activity of the neurons in the model and binning procedure over time windows of length *τ*. **c**) Point cloud representation of the binned spike train in, neural space with coordinates as activity of single neurons. **d**) Example of a symmetric distribution of activities for three neurons, while the rest are silent. **e**) Example of an asymmetric distribution of the activities. **f**) Dimensionality of the neural activities as a function of average connectivity in SONET networks, with varied average connectivity and motif statistics.

The stochastic spiking dynamics induced by Eq. 1 leads (cfr. Supp. Mat.) to an equation for the covariance matrix ***C*** of the network response. For simplicity we present the result as a matrix of spike train auto- and cross-spectra at frequency *ω*, ***C***(*ω*). This is the matrix of the Fourier transforms of the familiar auto- and cross-covariance functions; its zero mode ***C***(0) is the the usual covariance matrix on which we will focus for the rest of this work [47, 48]. Very usefully, this mode has been shown to yield an accurate approximation of correlations over any time window that is long enough to encompass the structure of neural correlograms [49]. The linearized dynamics, Eq. 1 give rise to the covariance matrix as:

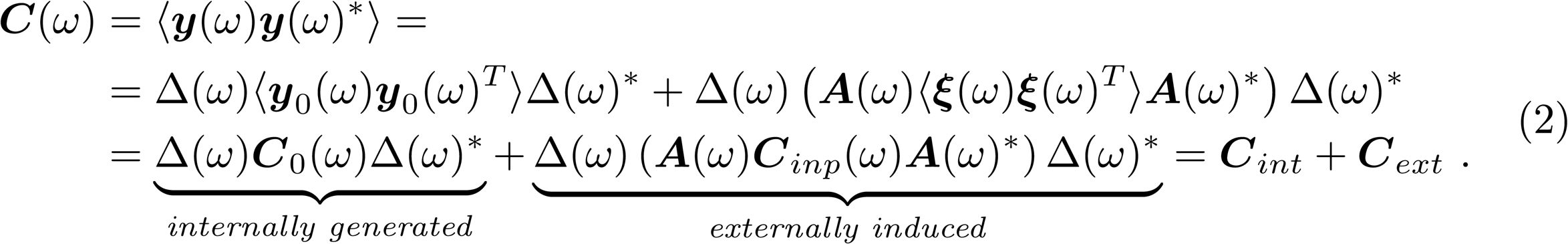

The first term of Eq. 2 expresses how the variability in the activity of single neurons (the baseline covariance ***C***_0_) propagates through the network to induce internally-generated covariability. Similarly, external inputs with covariance ***C*_*inp*_** give rise to covariances ((***A***(*ω*)***C*_*inp*_**(*ω*)***A***(*ω*)*^∗^*) in the externally induced term), which then propagate through the network. (External inputs with low-rank correlations could reflect global fluctuations due to shifts in attention, vigilance state, or motor activity [50, 38].)

Above we also introduced Δ(*ω*) = (***ℐ*** - ***G***(*ω*))^-1^, where Δ_*ij*_ is called a propagator as it reflects how a spike in neuron *j* propagates through the network to affect the activity of neuron *i*. Eq. 2 has been extensively studied in a number of frameworks [51, 52, 53, 54, 44, 55].

### 2.2 Measuring dimensionality

We aim to characterize the dimensionality of the distribution of population vector responses. Across many trials, these population vectors populate a cloud of points. The dimensionality is a weighted measure of the number of axes explored by that cloud:

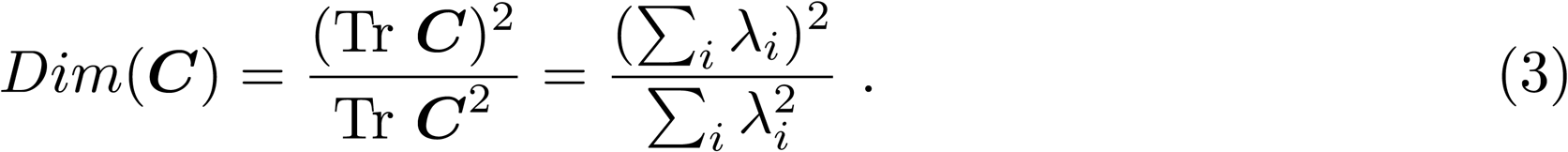

where *λ_i_* is the *i*^*th*^ eigenvalue of the covariance matrix ***C***. The eigenvectors of the covariance matrix **C** are the axes of such cloud of points as in Fig. 1c. If the components of ***y*** are independent and have equal variance, all the eigenvalues of the covariance matrix have the same value and *Dim*(***C***) = *N*. Alternatively, if the components are correlated so that the variance is evenly spread across M dimensions, only M eigenvalues would be nonzero and *Dim*(***C***) = *M* (Fig. 1d). For other correlation structures, this measure interpolates between these two regimes (Fig. 1e) and, as a rule of thumb, the dimensionality can be thought as corresponding to the number of dimensions required to explain about 80% of the total population variance in many settings [26, 3, 25].

Previous works have shown that the average correlation between neurons depends strongly on the motif structure of their connectivity [17, 18, 20]. We began by asking whether the same is true for the dimensionality. To do this, we generated random networks with a range of connection probabilities and, for each connection probability, a wide range of two-synapse motif frequencies (SONETs; Methods c and e, and [40, 56]). In Fig. 1d we plot the dimensionality of the network’s activity against the average probability of connection *p* (0 *≤ p ≤* 1) for an ensemble of SONET networks (cfr. Methods e for network details). The first notable observation from Fig. 1f is that the dimensionality for such networks is strongly influenced by *p*: as *p* increases, the dimensionality decreases towards 1. Importantly, Fig. 1f also shows a high range of variability in the dimension produced by networks with the *same* value of average connectivity *p*, indicating that the way that a given number of connections is arranged across the network also plays a strong role in determining the dimension of its activity Fig. 1f. Our next major goal is to describe how the statistics of connectivity motifs gives rise to this variability.

### 2.3 Expressing the covariance in terms of network motifs

We review the main ideas of the theoretical framework that allows for an expansion of Eq. 2 in terms of connectivity motifs. For a more comprehensive description see Suppl. Mat. and [17, 18]. This framework aims to model the complexity of connectivity structures in real world networks, like the one represented in Fig. 2a, in terms of motif statistics. There are three main conceptual steps to highlight. We will introduce them in the case where the network does not receive any external input so that ***C***(*ω*) = Δ(*ω*)***C***_0_(*ω*)Δ(*ω*)*∗* but they can be extended (cfr. Sec. 2.5 and Suppl. Mat. Sec. 2 to the more general case where such an input is present.

**Fig 2.**
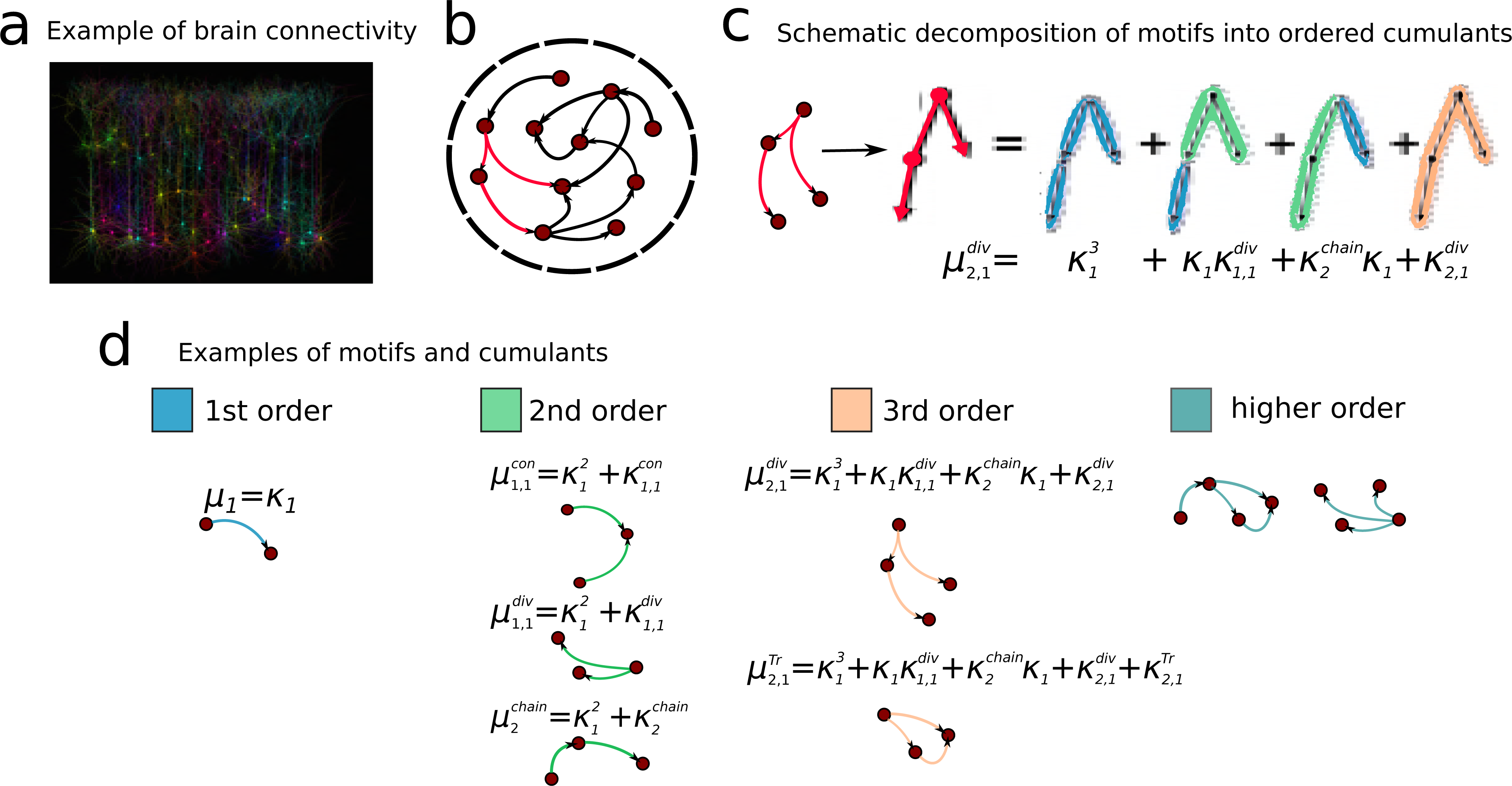
Description of connectivity motifs and cumulants. **a**) Example of brain cortical connectivity [57]. **b**) Motifs in the model of a recurrent neural network. In red is highlighted an example of a divergent motif. **c**) Example of the decomposition of a divergent motif into cumulants. **d**) Categorization of connectivity motifs into orders, with several example of the cumulants decomposition. Notably we highlight a novel motif: the trace motif *µ*^Tr^, as an example of a 3rd order motif.

The first step is to expand the propagator:

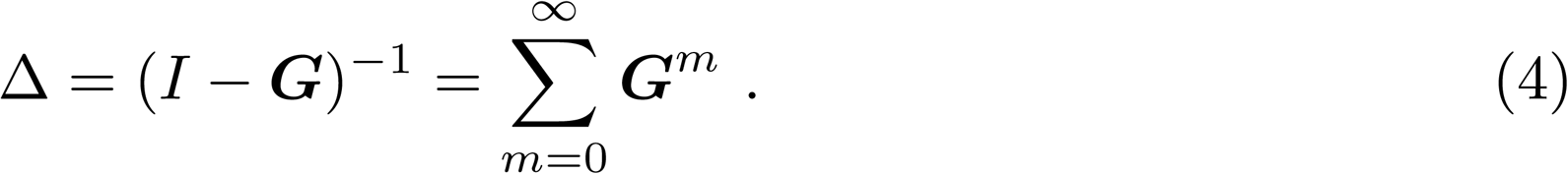

By expressing Δ in this form we can then write ***C*** (dropping the dependency on *w*) via an expansion:

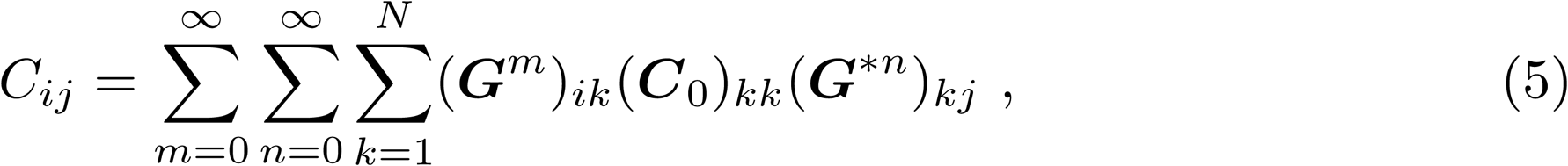

where from now on we will consider the case where ***C***_0_ is diagonal ***C***_0_ = *c*_0_ ***ℐ*** ‐ as for the standard assumption and model of initially independent Poisson neurons that are then coupled together into a network. Then Eq. 5 provides an intuitive description of the spike train cross-spectra in terms of paths through the network. This captures contributions to the cross-spectrum for paths that fork out of neuron *k* and end on one side in neuron *i* after *m* connections, and on the other side in neuron *j* after *n* connections. An example of such a path for *m* = 2 and *n* = 1 is shown in red in fig. 2b. The expression in eq. 5 has been studied extensively in previous works [58, 52, 53, 54, 20].

The framework in which we cast our theory relies on a second conceptual step, based on rewriting a function of the covariance ***C***, Eq. 5, in terms of motifs. In the case of the where this function is the average covariance 〈***C***〉, this takes the form [17]:

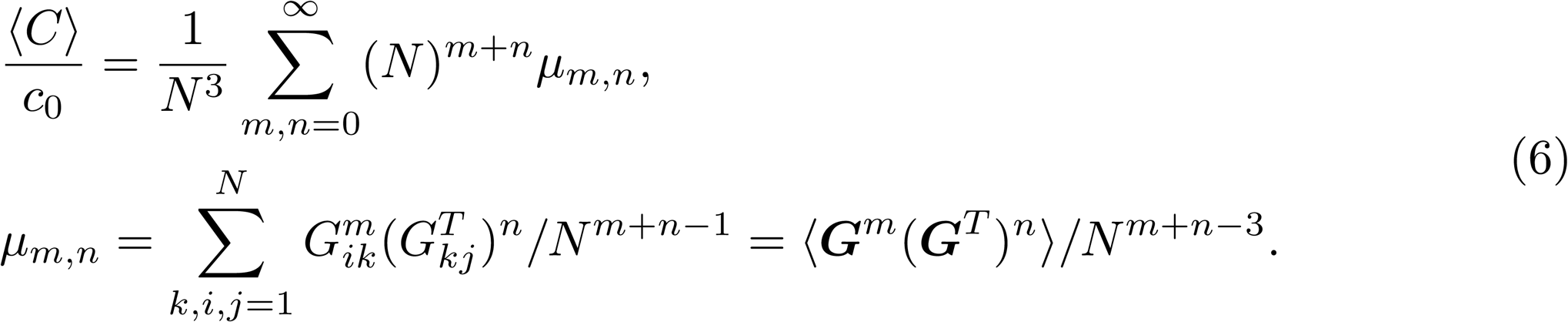

Here, we assumed that cellular response properties are homogeneous ***A*** = *g****ℐ***, and ***C***_0_ = *c*_0_ ***ℐ***. The motif moment *µ_m,n_* measures the average strength of a (*m, n*)-motif composed of two paths (respectively of length *m* and *n*) connecting any neuron k with neuron i and j. Examples of a (1, 1)-, and (2, 1)-motif are shown in Figs. 2c and 2d. Motifs of this kind, where paths originate from a common neuron, are called divergent motifs. We consider five kinds of motifs: convergent, divergent, chain, reciprocal and trace, depending on the direction of edges to the common node as illustrated in Fig. 2d. These motifs correspond to similar definitions to the one for *µ_m,n_* in Eq. 6 (cfr. Suppl. Mat. sec. 2.1 for additional details). In networks where all synaptic weights have same value, then *µ_m,n_* is proportional to the frequency of the motif.

We can also define weighted motif statistics. For example:

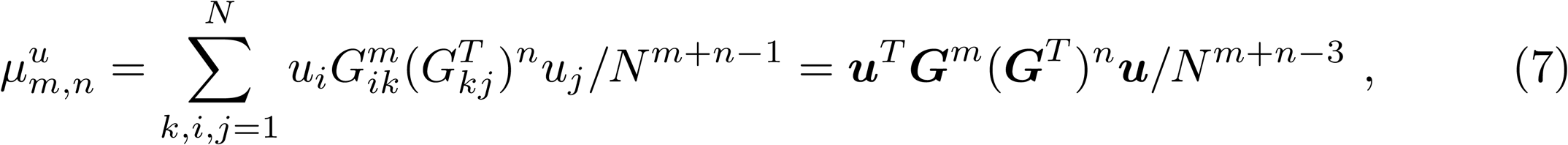

where ***u*** is a vector of norm 1 (*||u||* = 1). For example, ***u*** could contain neuron’s firing rates, or be the eigenvectors of ***W***. The case of Eq 6 corresponds to choosing the unit norm vector of constant entries, 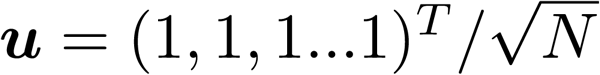. Ultimately the choice of ***u*** depends on the desired function of the covariance to compute (e.g. ⟨ ***C*** *⟩*, Tr(***C***), *Dim*(***C***)*…*), on the structure of ***G***, and on the presence or absence of inputs. In what follows this choice will be motivated in each case.

The last and crucial conceptual step of the theoretical framework is to re-sum the motif moments by rewriting them in terms of cumulants. The idea is to approximate the probability of finding a specific motif *µ_n,m_* by iterative approximations built through the probabilities of finding the building blocks of that motif. For example, in Fig. 2c we see how the probability of motif *µ*_1,2_ to occur in the network can be subdivided in the probabilities of finding its building blocks: three synapses 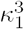, one synapse and one chain of length two *κ*_1_*κ*_2_ and so on. The general relationship between moments and cumulants is [18]:

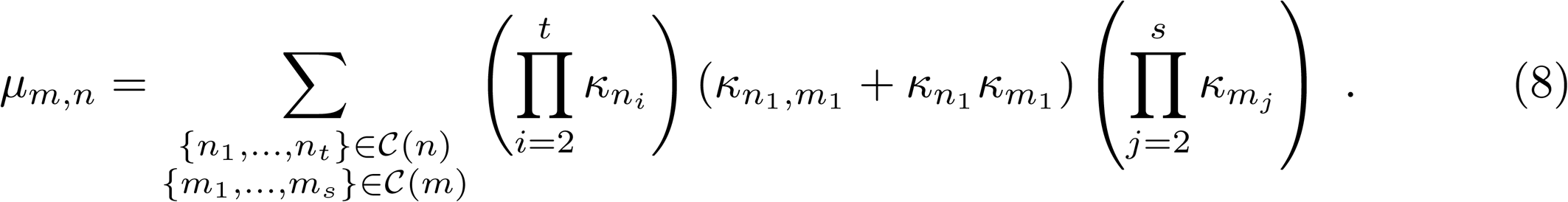

where each *κ_n_, κ_n,m_* is a cumulant (respectively for chains and divergent motifs) and *C* (*n*) is the collection of ordered sets whose elements sum up to *n*. This step removes redundancies and improves the rate of convergence of the expansion, so that only relatively smaller motifs need to be measured and included. This is accomplished by “resumming,” via the identity:

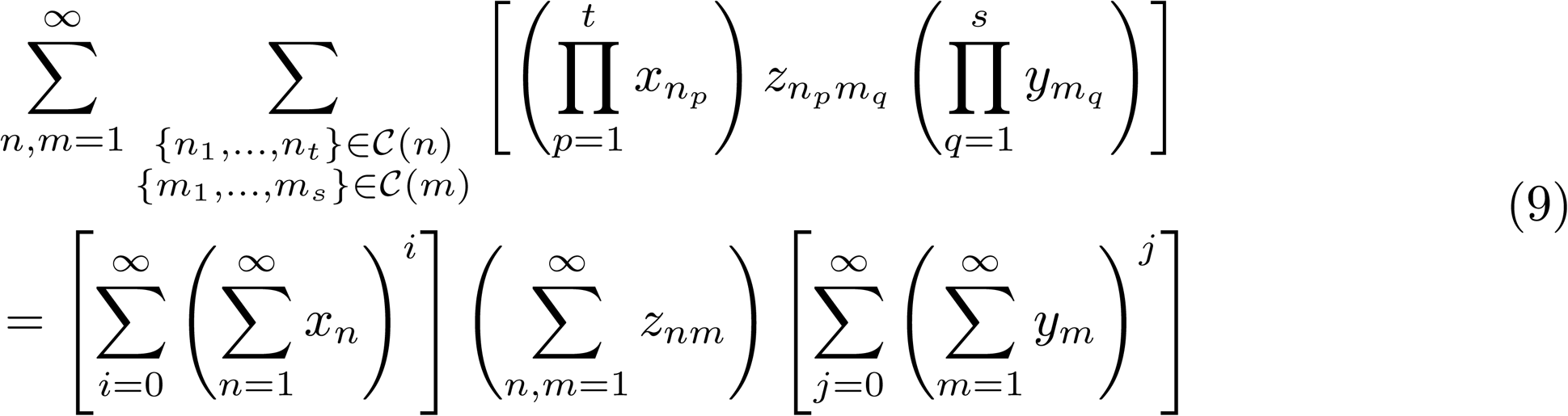

that allows one to resum the contribution of each cumulant to any order in the expansion of Eq. 5. In this way the expression for a function of the covariance matrix assumes a closed form as a function of the cumulants (e.g. Eq. 6 for the mean covariance).

Through the resumming procedure we are computing the contribution of any cumulant *κ* not to a specific term ***G***^*m*^(***G***^*T*^)^*n*^ but to the full sum 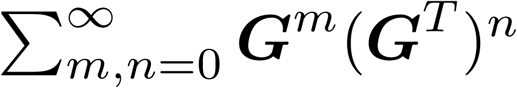. In summary, this approach allows us to remove redundancies in motif statistics, and to isolate the impact solely due to higher order motif structures [17, 18].

The framework outlined above results in the ability to write any function of the covariance in terms of motif cumulants. Specifically, according to our interest here, the expressions for ⟨***C*** ⟩ and *Dim*(***C***) can be written in terms of a small subset of cumulants. In the following (cfr. Suppl. Mat. sec. 2.4) we will explain how this framework can be deployed in computing *Dim*(***C***) for different networks, first in the absence of inputs, and then in their presence.

### 2.4 Dimensionality of internally generated network activity

In our results we will include cumulants up to second order, although the expansion and theory can be taken to higher order. Second order cumulants correspond to chains 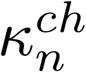, convergent paths 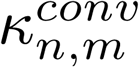, divergent paths 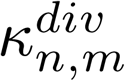, reciprocal paths 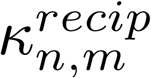 and trace motifs 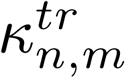 as shown in Fig. 2d. Mathematical definitions and more detailed explanations of the meaning of these cumulants can be found in the Suppl.Mat. sec. 2.1-2.4.

The expansion in terms of cumulants leads to the expression for the average covariance ⟨***C*** ⟩ ([18]):

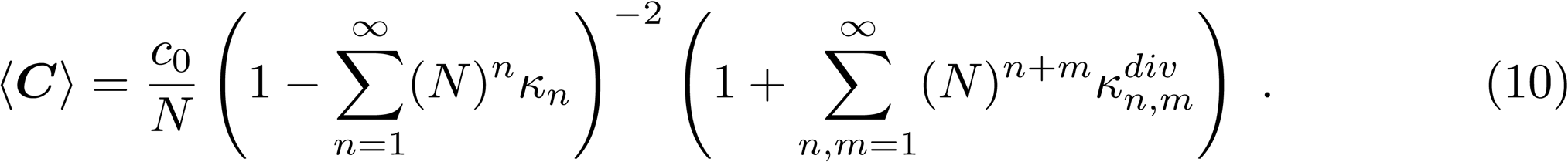

Notably, the contributions of chains and divergent motifs factor out in Eq. 10.

The expression for the dimensionality *Dim*(***C***) is the ratio between Tr(***C***)^2^ and Tr(***C***^2^), and these two quantities are general functions of the cumulants so that

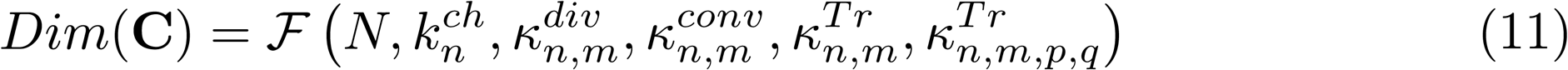

where ℱ is a function whose full expression is shown in Methods a, in terms of its numerator Tr(***C***)^2^ and denominator Tr(***C***^2^). This full expression also shows that the dimensionality is directly related to the average covariance ⟨ ***C*** ⟩. Specifically, it turns out that the dependency of *Dim*(***C***) on 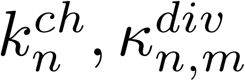 is the same as that of ⟨ ***C***⟩, so that we can rewrite Eq. 11 as:

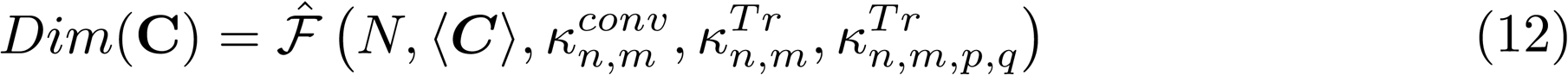

highlighting the role of convergent and trace motifs in regulating the relation between the average covariance and the dimensionality (a detailed expression of Eq. 12 can be found in Suppl. Mat. sec. 2.3). The trace cumulants 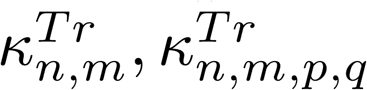 in Eq. 11 represent the statistics of motifs corresponding to patterns of connectivity that originate in one neuron and converge to a second neuron (Fig. 2d). We will show later how these statistics are highly correlated with reciprocal connections.

We next interpret and apply the formulas just described, which predict the dimension of network-wide activity in terms of localized connectivity motifs. We first use two classes of networks as examples: “purely random” Erdos-Reyni networks, and an exponential family of random graphs parameterized by second order motif statistics. While these are quite natural (but by no means automatic) cases for our theory, which is based on localized connectivity statistics, to succeed, we later apply it to different types of complex networks.

We begin by analyzing an interesting limit of Eq. 11: an Erdos-Reyni network. For a Erdos Reyni network all cumulants except for 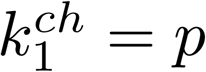 (where *p* is the probability for each edge to be present in the graph) and the trace cumulants

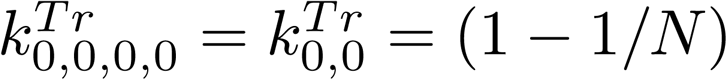 are zero. In this limit Eq. 3 becomes:

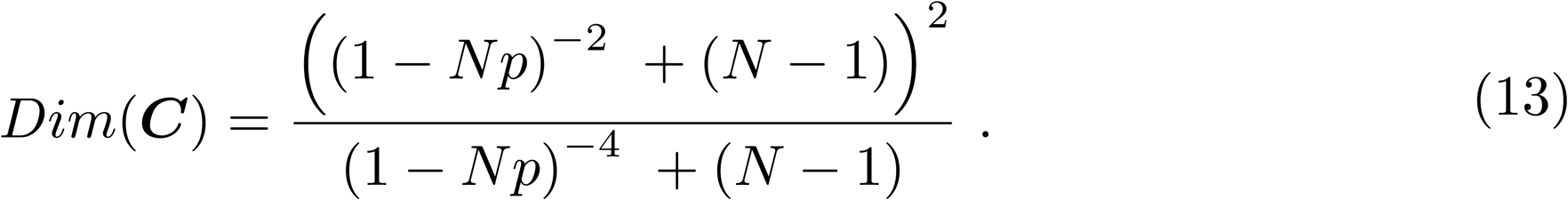

From this expression we see that when 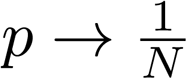 we obtain *Dim*(***C***) *→ N -* 1.

This behavior can be interpreted in the following way: for *p* small enough that the structure of ***C*** is fully diagonal and all the elements are equal to *c*_0_; in this regime all the neurons in the network act independently and contribute equally to *Dim*(***C***). As *p* increases more and more neurons start interacting and the dimensionality decreases until we obtain *Dim*(***C***) = 1. In Fig. 3a we see how Eq. 13 (red dashed line) is in agreement with the full expression for *Dim*(***C***) (green line) where Eq. 2 has been used for the internally generated covariance in spite of the cumulant approximation.

**Fig 3.**
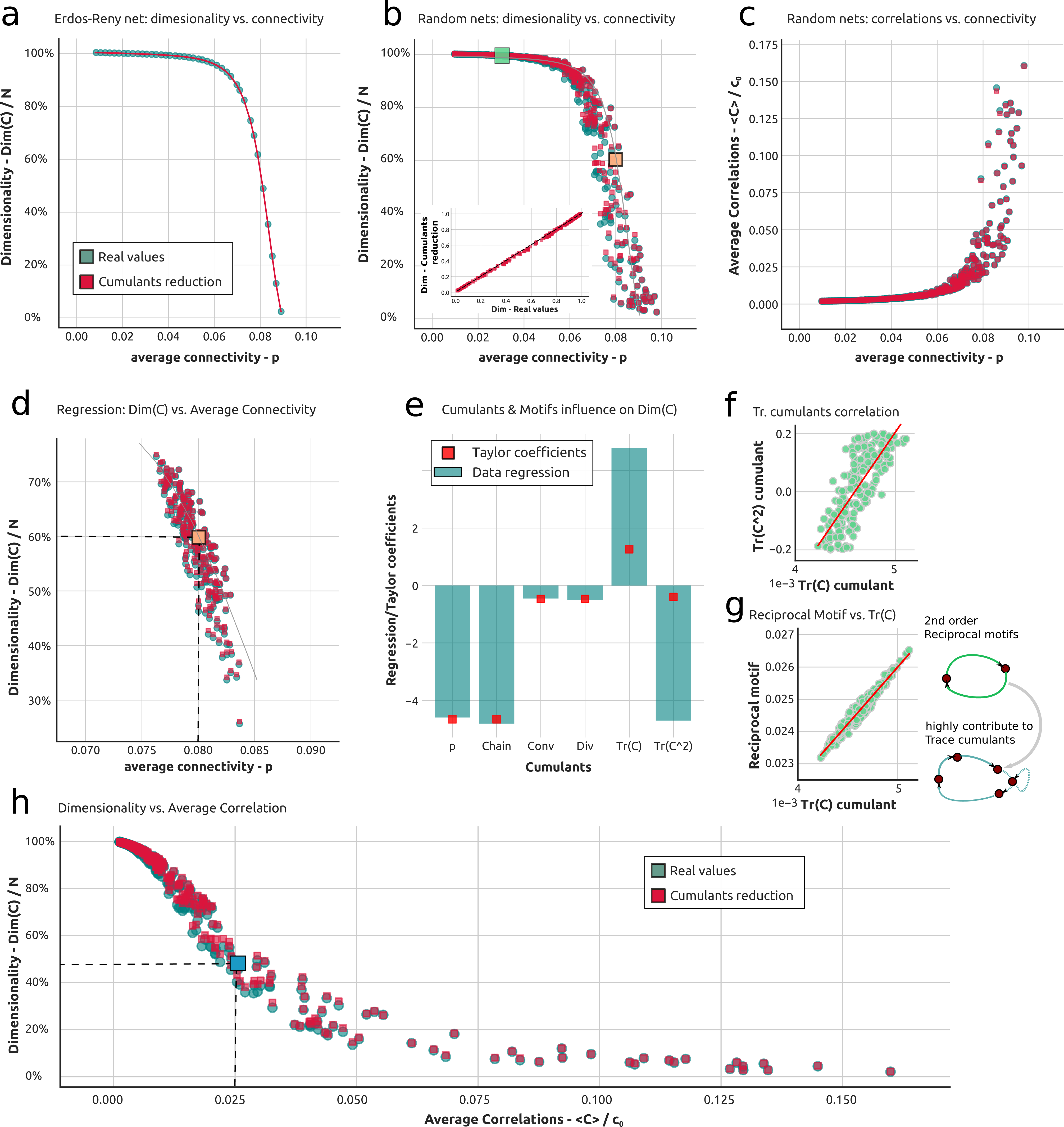
Theory of dimensionality in random excitatory recurrent networks through connectivity motifs. **a**) Dimensionality as a function of average connectivity in an Erdos-Renyi network. The full theory is in green while the theoretical approximation via the cumulant framework is shown in red. This color code is consistently used throughout the figure and paper. **b**) Dimensionality as a function of average connectivity in SONET networks. Highlighted in green (orange) is a point corresponding to a weakly (strongly) connected network. The inset shows the exact value of the dimensionality versus the approximated one. The gray line follows the ER case of panel a). **c**) Average correlation vs. average connectivity in SONET networks. **d**) Dimensionality vs. average connectivity in the ensemble of SONET networks used for the regression. Highlighted in orange is the point corresponding to the Erdos-Renyi network where the Taylor expansion is centered. **e**) Comparison between regression and Taylor coefficients. In green are the regressors of the multilinear regression of the dimensionality regressed against the cumulants, while in red are the Taylor coefficients of the expansion around the Erdos-Renyi network highlighted in panel e. **f**) Relation between trace cumulants for the ensemble of networks used in the regression. **g**) Relation between the trace cumulant and the reciprocal motifs in the ensemble of networks used in the regression. **h**) Dimensionality versus average correlation for the ensemble of networks used for panels b and c. The blue point highlights a point with relatively low dimensionality for a relatively low (and commonly observed) value of the average correlation.

To show the efficacy of Eq. 11 in capturing the dimensionality of network responses, we use this expression to compute *Dim*(***C***) in an ensemble of SONET networks [40]. These (cfr. Methods f) are random networks where the probability of having a second order motif can be arbitrarily modified; such networks can therefore assume a wide range of values for second order motifs and cumulants. In Figs. 3b and 3c we show the dimensionality and average correlation values (given by ⟨ ***C*** ⟩ */c*_0_) for a wide range of SONET networks, with a network’s dimensionality plotted against its connection probability *p*. Here, for each network we plot both the dimension computed via the full covariance formula Eq. 3, as well as via the cumulant truncation via Eq. 11 (red dots). Although the dimensionality varies strongly across networks with different motif statistics even at a fixed value of *p* (as was already pointed out in Fig. 1f), the cumulant theory matches this variability closely across the range of SONET networks. This is shown in Fig. 3b in two ways: for each network (every green dot) the corresponding theoretical approximation (corresponding red dot) lies right on top or closeby; the inset in Fig. 3b confirms this by plotting dimension calculated via the cumulant approximation against the true values from the full covariance expression.

Together Fig. 3b shows that second order motifs contribute to the dimensionality of the response according to Eq. 11. However, from Fig. 3b it is not possible to single out the contribution of each motif. To address this question we consider an ensemble of SONET networks centered in their statistics around an Erdos-Renyi network with *p* = 0.08, corresponding to the orange dot in Figs. 3b and 3d (see Methods f for details). The dimensionality for the response of each network in this ensemble is plotted against *p* in Fig. 3d. Then we carry out a multilinear regression (see Methods f) of the dimensionality of this ensemble of networks against the values of each cumulant. The regression coefficients express how each cumulant influences the dimensionality (*r*^2^ = 0.994) (Fig. 3e) so that:

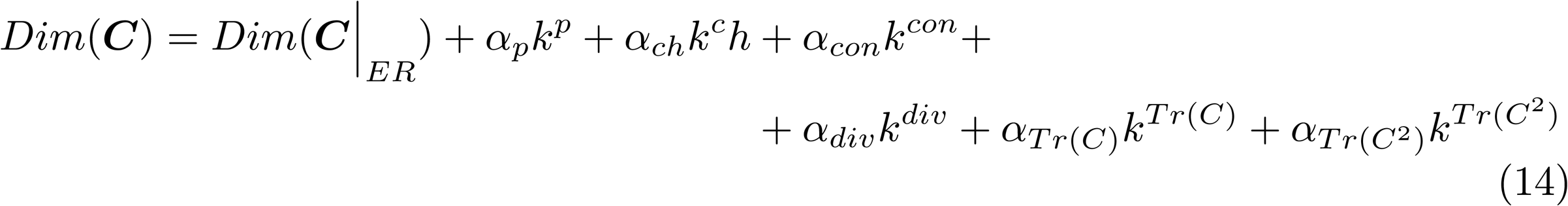

where the *α*^/′^*s* are the regression coefficients for each cumulant (green bars in Fig. 3d). An increase of most cumulant, but not all, types of cumulants appears to lead to a decrease in dimensionality as most coefficients in Fig. 3e are negative. This is important as it suggests that adding most types of connectivity structure to a circuit generally lowers the dimensionality of the response.

In more detail, this analysis shows that, while increasing the average connectivity, chains, and diverging and converging motifs leads to a decrease in dimensionality, terms contributing to the trace motifs may play a role in expanding the dimensionality. Complicating matters is that 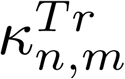 and 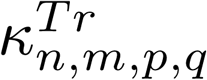 are, in general, highly correlated in their values. This correlation is shown in Fig. 3f and it limits the applicability of the regression to the ensemble with respect to the trace cumulants, as can be seen in Fig. 3e. To get a theoretical handle on this, we analytically compute the Taylor coefficients of the expansion of the dimensionality formula Eq. 11 in terms of motifs, expanded around the Erdos-Renyi case (orange point in Fig. 3d). The Taylor coefficients are the *α*^*′*^*s* in the first order theoretical expansion of Eq. 14 of the dimensionality formula. To ease reading of the resulting formulas we first define:

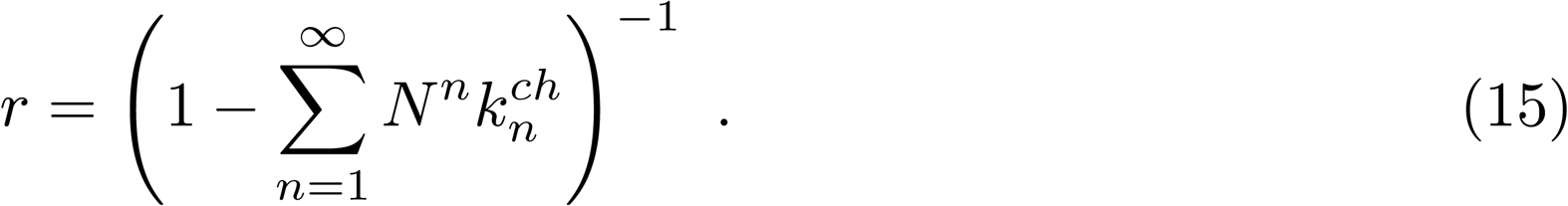

The expressions for Tr(***C***) and Tr(***C***^2^) in the Erdos-Renyi case have then the form:

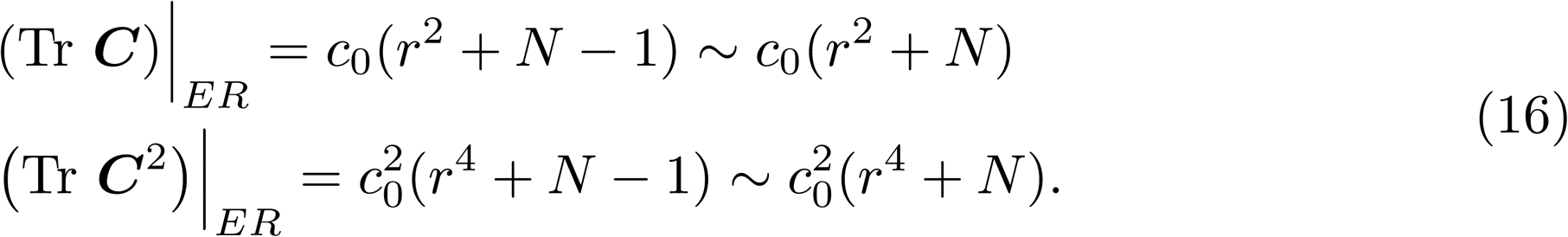

The expressions for the Taylor coefficients of second order motifs are:

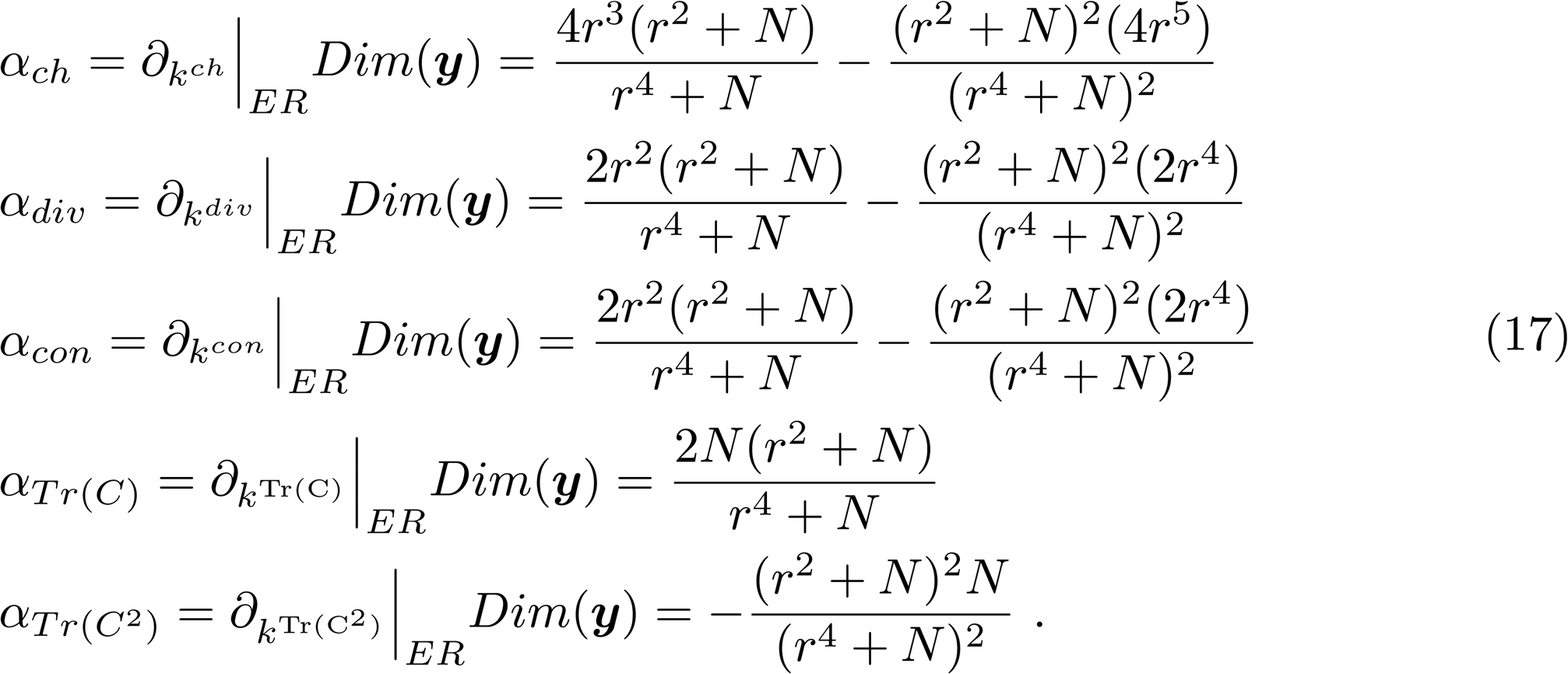

A derivation with more details is available in the Suppl. Mat. 2.5. These expressions represent the corresponding theoretical quantities for the regression coefficients of Fig. 3e and are shown as red dots. As we see the Taylor coefficients provide a direct understanding of the effect of increasing different cumulants on the dimensionality. Moreover, as we show analytically in the Supplemental material, *α_ch_ <* 0, and *α_div_* = *α_conv_ <* 0; thus, the effects of adding chain, diverging, or converging motifs to a given network is to drive down the dimension of the activity that it produces.

Although the regression fails to capture the right quantitative expressions for the trace motifs (see Fig. 3e), it does suggest that these terms play a key role in regulating the dimensionality. Trace cumulants are mainly influenced by reciprocal motifs as they directly enter the computation for the trace cumulants. This can be observed in Fig. 3g where the high correlation between the two is highlighted. Altogether these results point to reciprocal connections as major players in balancing the overall behavior of the dimensionality.

**Dimensionality versus average covariance** Finally, in Fig. 3h, we show how the dimensionality is related to the average pairwise spike count correlation across the range of SONET networks. Importantly, we see that dimensionality attains very low values, even when the average correlation values are very weak. For example, when average correlations ⟨***C***⟩/c_0_ = 0.025, we see that *Dim*(***C***) = 0.5 (blue point in Fig. 3h). In other words, when cells appear almost uncorrelated on average (*≈* 2 - 3%), the overall dimensionality of spiking activity can be cut by half compared with the uncoupled case.

While this phenomenon could be foreseen by looking closely at Figs. 3b and 3c we highlight it here as it helps to reconcile two observations often seen in the literature: relatively weak activity correlations [59, 34, 43] yet relatively low activity dimension [1]. We note that [43] has made closely related findings about highly restricted sets of firing patterns that can be implied by weak pairwise correlations. In our framework, the dimensionality can be tightly linked to the average covariance, see Eq. 12, but we also show that converging and trace motif cumulants influences the dimensionality but not the average covariance (see Eq. 12 and Eq. 22 in Methods a). This points to dimensionality not only as a comprehensive measure of how coordinated network activity is, but also also as a more sensitive means to assess how coupling is coordinating that network activity (cfr.[43]). We will further expand on this important point in the Discussion.

### 2.5 Dimensionality of stimulus-driven responses

In Eq. 2 we highlighted two contributions to the total covariance of the network activity. The first is due to the internally generated activity (the reverberation of the stochastic Poissonian spiking through the network), and the second is due to the inputs to the network. While in the previous sections we have analyzed the dimensionality of the network response in the absence of inputs, here we generalize the results to include their contribution. The interplay between the connectivity of the network and the inputs can be captured by Eq. 3 where we expressed ***C*** as ***C*** = ***C*_*int*_** + ***C*_*ext*_**:

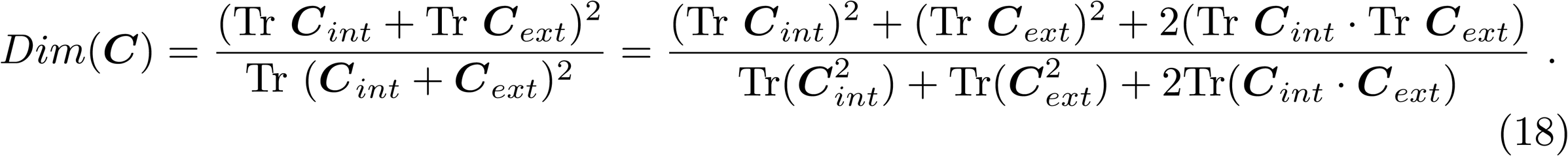

We decompose the input covariance ***C*_*inp*_ into *N*_*inp*_** orthogonal unitary factors ***ξ***, so that 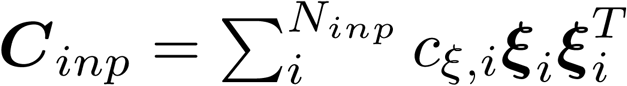. The external input to the network might arise from the spontaneous or evoked activity of other areas; regardless, it can be modeled as a sum of independent contributions where the number of factors *N*_*inp*_ and the individual strength of these factors *c****_ξ,i_*** has to be determined.

The theory introduced in Sec. 2.3 needs to be extended to reflect a crucial fact: the input may target different neurons in the network to a different degree. In turn, connections from and to specific neurons will be more important than others in driving network-wide activity. In Sec. 2.3 and Fig. 2, we introduced motif moments and cumulants by specifying that weights from different neurons were equally taking part to the computation of the dimensionality. Thi*v*s <u>id</u>ea was rendered mathematically by using a uniform weight vector 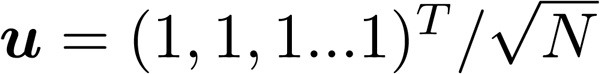 in defining and resumming motif cumulants. In the following we will also employ a set *N*_*inp*_ of vectors ***u****_ξ,i_* = ***ξ_i_*** to properly resum different contributions to the input structure and their reverberation through the network. In Suppl. Mat. sec. 3.2 we show how all these contributions can be dealt with and re-summed simultaneously via proper handling of *weighted* motifs and cumulants, building from [17, 18]. Here, each motif simply carries a weight corresponding to the product of input strengths for each of the neurons that compose it.

The resulting equations have function forms similar to the one of Eq. 11, but with weighted cumulants. Denoting with *κ_ext_* the set of input weighted cumulants and with ***κ*_*int*_** the set of internal cumulants employed in Eq. 11, we have:

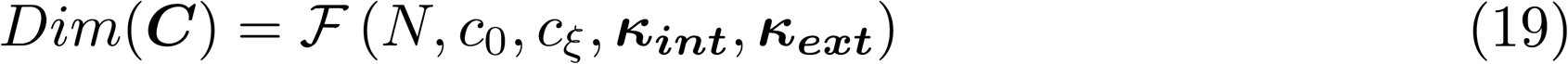

The full expression for this equation, in terms of its building blocks of Eq. 18, is given in Methods b. This equation formalizes the interplay between stimuli, connectivity, and internally generated activity in creating network activity with a particular dimension. In what follows, we illustrate one aspect of this: how the strength and dimensionality of inputs to a network modify the “total” dimensionality of the network responses. While the limiting trends are exactly what one would expect – stronger inputs increasingly entrain the network response, and higher dimensional inputs lead to higher dimensional responses – both the limiting values of dimensionality and the approach to them depend on details of network connectivity.

To better illustrate this process, we study the response of two different networks: a weakly and a strongly connected network. These two cases correspond to the two points highlighted in Fig. 3b: the green point (*p* = 0.03) to a weakly connected network, while the orange one (*p* = 0.08) to a more strongly connected one. In both cases the internally generated activity is uniformly weak or strong across all neurons. To gain more insight on how skewed distributions of intrinsic variances would affect our analysis we refer the reader to [25].

To begin, consider a weakly connected random network receiving *N*_*inp*_ input factors, each with the same strength *c****_ξ_***, so that 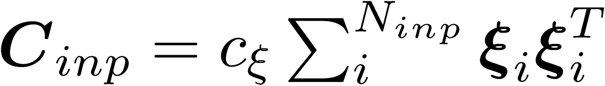 We examine the dimensionality of the network response as a function of *N*_*inp*_ in Fig. 4a. Note that as *N*_*inp*_ grows, *both* the dimension of the input (*N*_*inp*_) and its overall strength (variance N_*inp*_c_ξ_ grow. The initial dimensionality in the absence of any input is close to 100%, then it decreases as more and more inputs are fed into the network, eventually growing with the number of inputs as these entrain the network activity. Both the extremes have dimensionality close to 100%, as shown in Fig. 4a, and in between there is a trade-off region where the low dimensionality of the input and the high dimensionality of the internal activity interact non-linearly as shown in Eq. 18. To better understand these trends we rewrite Eq. 18 by using Eq. 2 with ***C***_0_ = *c*_0_***ℐ***, ***A*** = *g****ℐ*** and 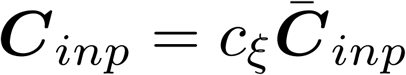 where we have highlighted the scaling factor of ***C*_*inp*_**. The resulting expression is:

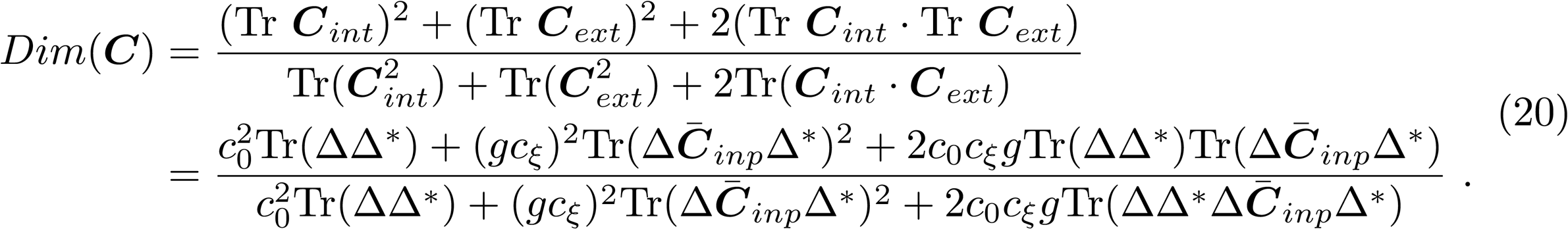

**Fig 4.**
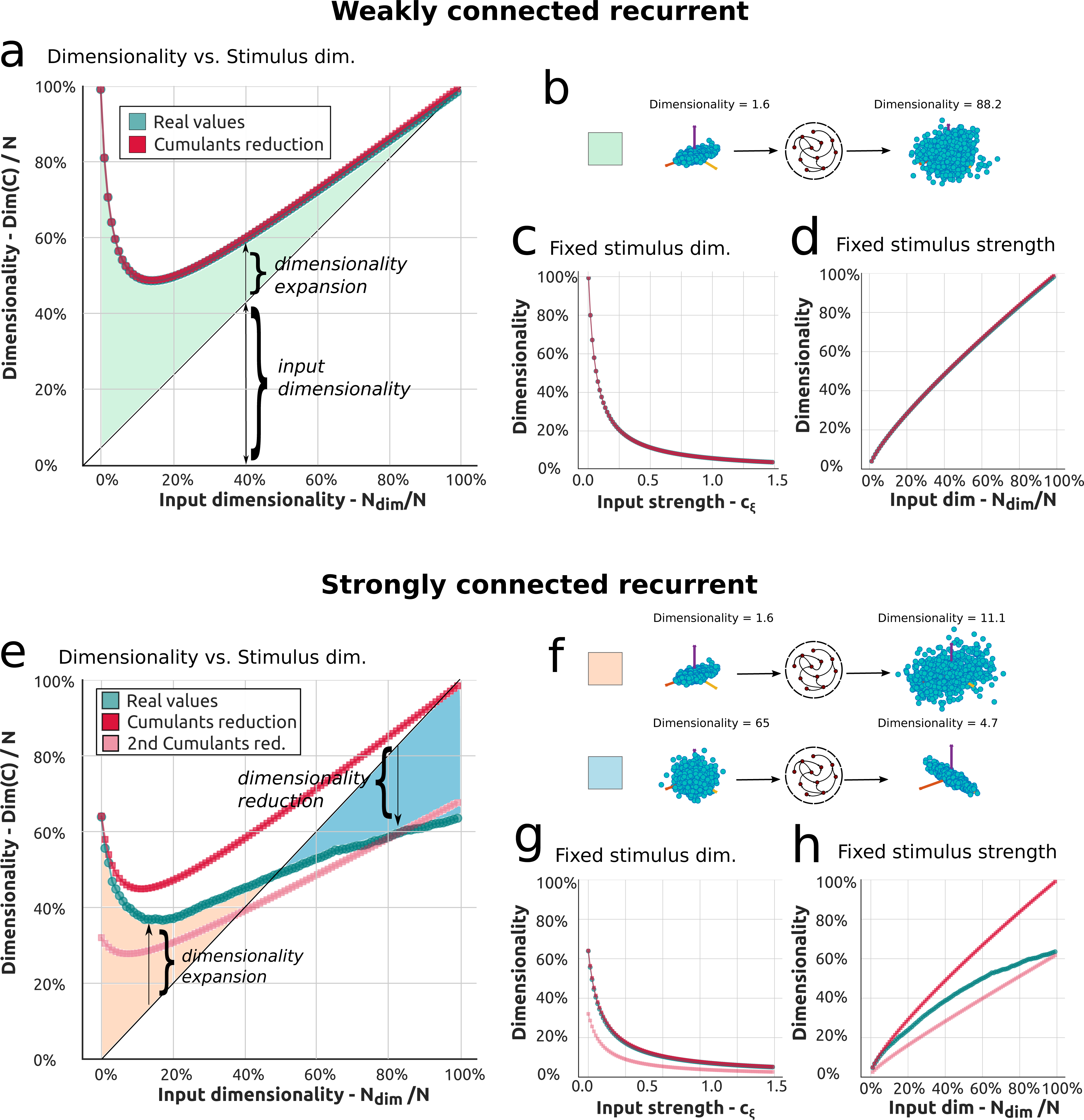
Dimensionality of the network response in weakly and strongly connected excitatory recurrent networks. **a**) Dimensionality of stimulus driven responses as a function of the dimensionality of the stimulus in a weakly recurrent network (see text for important details on how the stimulus is defined). The line in green is the full theory while the line in red is the theoretical approximation in the cumulant framework. In light green is the area that marks the region of expansion of the dimensionality with respect to the input. **b**) Example of the expansion of the input to the network, schematized by the effect of the network in inflating the cloud of points. **c**) Dimensionality versus stimulus strength for a unidimensional input. **d**) Dimensionality versus stimulus dimensionality for a stimulus of fixed strength. The total strength is rather high so that the initial dimensionality for a unidimensional input is extremely low. **e**) Dimensionality of stimulus driven responses as a function of the dimensionality of the stimulus in a strongly recurrent network. The line in green is the full theory while the line in red is the theoretical approximation in the cumulant framework. In pink is a second approximation in the cumulant framework that accounts for a high dimensional input. The areas in orange and blue are mark respectively the cases of dimensionality expansion and reduction. **f**) Cartoons for examples of dimensionality reduction and expansion induced by the internal modes of a strongly recurrent network. These behaviors are induced by the strongly recurrent connectivity. **g**,**h**) Analogous panels to panels c and d for the strongly connected case.

In this formula we recognize that the limits highlighted above (absence of input regime and input dominated regime) correspond to the cases where either the terms in 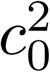 or in 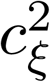 dominate, while intermediate cases are trading-off the contribution due to internal dynamics or external input. The distance of the full dimensionality from the diagonal (green region) measures the dimensionality expansion, where the input distribution is “inflated’ by the network’s noisy internal dynamics, Fig. 4b.

The non-monotonic behavior displayed in Fig. 4a can be explained as a trade-off of the two input properties introduced above: the input strength and dimensionality. The effect of the former can be understood in Fig. 4c, where we show how the dimensionality of the response decreases as a function of a gradually stronger unidimensional input (*N*_*inp*_ = 1 and increasing *c****_ξ_***). This behavior can be compared to established properties of stimulus driven dynamics in cortical circuits [25, 60] where it has been observed that evoked activity suppresses the dimensionality of spontaneous activity. The influence of the latter factor, input dimensionality, is displayed in Fig. 4d where we provide the network an input of overall constant strength, of standard deviation 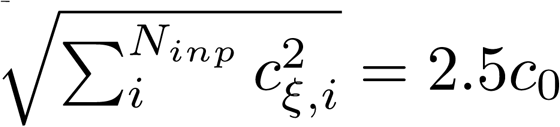 (cfr. Methods g), with increasing number of factors (dimensions). In this case, as the inputs fully entrain the network response, the dimensionality constantly increases. The trend in Fig. 4a can be interpreted as a trade-off between these two trends, again recalling that stimulus dimension and strength increase together in that plot Figs. 4c and 4d.

If we describe Fig. 4a as passing stimuli into weakly coupled networks leading to an expansion of the input dimensionality, then fig:4e shows that strongly coupled networks leads to a more complex trend. At first the input dimensionality is expanded, but then it is compressed; overall, the network response never achieves the full dimensionality of the input. In other words, the response is always constrained by the network dynamics: a first phase of dimensionality expansion is followed by a second phase of dimensionality reduction (Fig. 4f).

These two phases can both be understood qualitatively in terms of the propagator Δ in Eq. 2, that restrains the total network dynamics. In Fig. 4e the theoretical prediction made by the second order cumulant framework (red line) agrees with the exact dimensionality from formula Eq. 18 (green line) only for a low dimensional input, but then departs. This can be attributed to the many ways with which the inputs can interact with the internal modes of the network: as the number of input factors increases, evidently, the term ***C*_*int*_**·*C*_*ext*_ in Eq. 20 can no longer be captured by low order motif cumulants. In particular the motif cumulant approximation tends to overweight the importance of the input: the predictions for high *N*_*inp*_ in both Figs. 4a and 4e are similar. To weaken this limitation we show (pink line in Fig. 4e) a second theoretical approximation, where the terms arising from the internal modes are disengaged from the input contribution in Eq. 18. See Methods g for more details. This approximation captures more closely the properties of the network when, in the case of high dimensional input *N*_*dim*_ the activity is mainly constrained by the internal modes. We denote the two approximations, red and pink lines in Figs. 4e to 4h, respectively as the low and high dimensional input approximation. These two limits taken together show how low order cumulants are able to predict general trends in the dimensionality of driven responses.

Altogether we have shown in Fig. 4 how the interaction between the input and the network dynamics gives rise to a number of scenarios where the input dimensionality can be expanded, reduced or somehow controlled through the internal recurrent dynamics. Specifically we point out three different scenarios:

- If the input has low dimensionality and the network has high dimensionality due to weakly recurrent connectivity, the network expands the dimensionality of the input. The dimensionality expansion is effectively an “inflation” of the input dimensionality into the high dimensional neural space of the recurrent network (see Fig. 4b).
- If the input has low dimensionality and the network has also a low dimensional internal response due to strongly recurrent connectivity, the network still expands the input dimensionality. This mechanism (see Fig. 4f first case) is obtained as the input interacts with the internal activity of the network and their interaction adds up to create a new representation with higher dimensionality. This is mainly due to the constructive interaction between the internal and external covariance in the numerator of Eq. 18.
- If the input has high dimensionality and the network has low dimensionality due to strongly recurrent connectivity, the network reduces the input dimensionality. This results from a “bottleneck” induced by the low dimensional recurrent dynamics of the network (see Fig. 4f second case): the internal dynamics restrict high dimensional inputs to a lower dimensional subspace, as all they are projected onto the dominant eigenvectors of the network.

These points, as illustrated in Fig. 4, will be revisited in Sec. 3.

### 2.6 Complex and excitatory/inhibitory networks

Our results so far have shown a variety of phenomena in which the connectivity of a recurrent spiking network, and its resultant internal dynamics, shape its dimensionality. We have shown how this spectrum of behaviors can be interpreted in terms of the statistics of connectivity motifs: the theoretical framework introduced in Sec. 2.4 and illustrated in Figs. 3 and 4 points to motif cumulants as the logical building blocks. Moreover, truncating motif cumulant expansions at second order, so that only very localized connectivity data enters, can lead to quantitatively accurate predictions of dimension of intrinsic network activity and qualitative predictions of trends in the presence of stimulus drive.

This said, above we have tested these results only for fully excitatory random networks, and for those that are either fully random (Erdos-Reyni) or are generated according to low order connectivity statistics (SONET networks). It is possible that either the theoretical framework proposed (cfr. Fig. 3) or the dimensionality phenomena analyzed (cfr. Fig. 4), may not generalize to more complex networks. To attest this, in this section we generalize the results to complex networks with other structures, and with both inhibitory and excitatory neurons.

In Sec. 2.5 we introduced weighted cumulants to account for an input that was fed unevenly into different neurons within a networks. This is necessary as the cumulants originating in some neurons may have more impact on the network dynamics than others. The same argument holds true for the way internal activity in a recurrent network is generated intrinsically in a network, as some neurons, and their connectivity patterns, are known to have a stronger influence [61, 62]. Thus, we make use of generalized motifs for the internal activity of the network, in a similar fashion to what was done in Sec. 2.5 to account for input effects, by using weight vectors ***u*** in Eq. 7 that are chosen to be the eigenvectors of the connectivity matrix ***G***. This choice is justified by the same logic as in the case of the input: the directions identified by the eigenvectors are the ones where the activity propagates, so that neurons which participate more to the dynamical mode of the network are weighted more in computing the cumulants. Weighting neurons and thus motifs in such a way therefore handles the relative importance of cumulants in propagating activity through the network (see Suppl. Mat. 3.2 for further details).

To generalize our results we start by showing that the findings in Fig. 3 hold true for a wider class of complex networks. Specifically we compare three different network topologies: the Erdos-Renyi case studied before, together with Small World and Free Scale (Albert-Barabasi model) networks. For each case, we vary a single common parameter (cfr. Methods h), the density (probability) of synapses in the network *p*. In Figs. 5a to 5c we show three examples of the underlying weight matrices, one for each topology.

**Fig 5.**
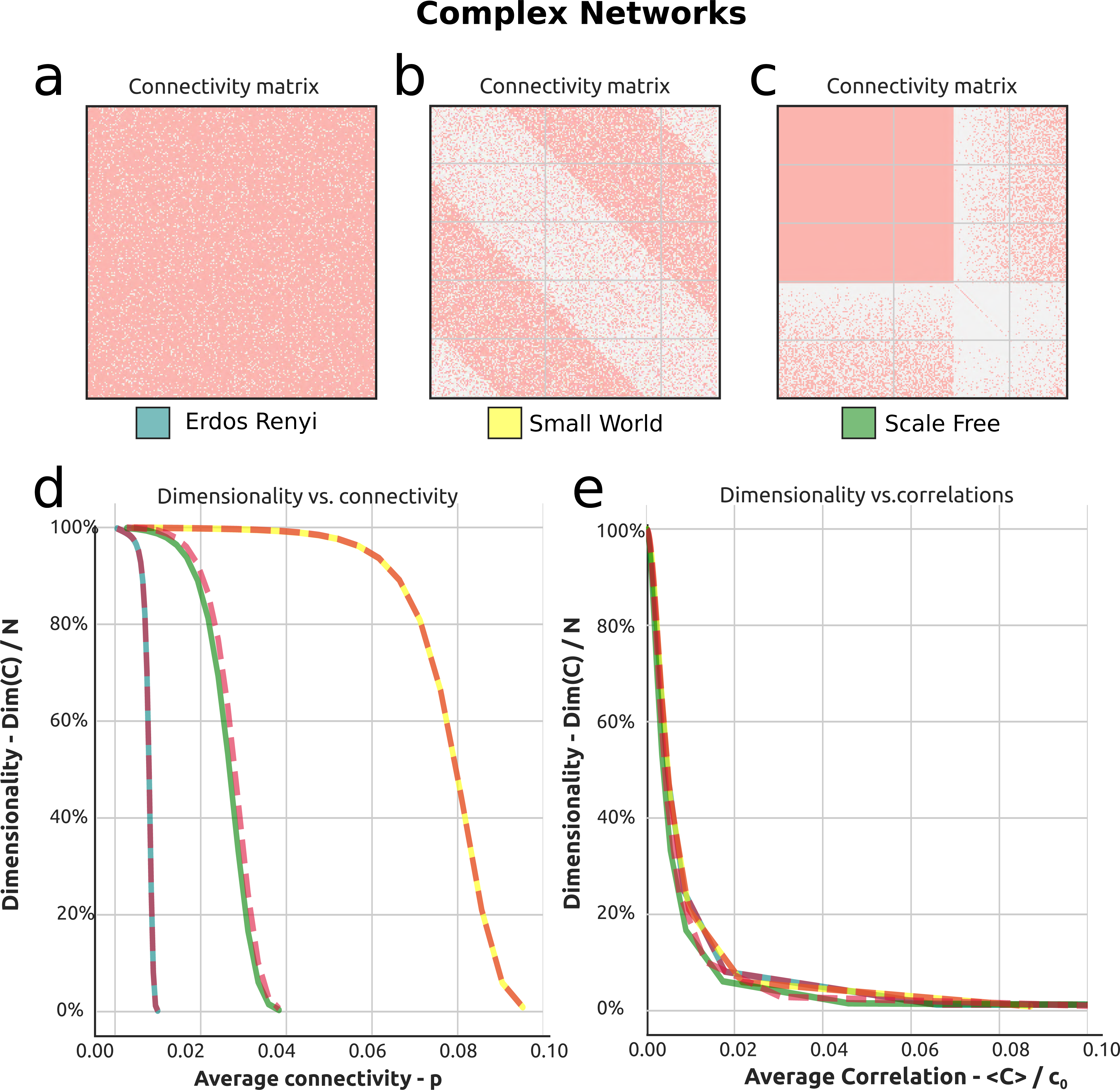
Dimensionality of networks with complex excitatory connectivity. **a,b,c**) Connectivity matrix ***W*** for the networks considered here: respectively Erdos-Renyi, Scale-Free, Small-World. **d**) Dimensionality as a function of average connectivity for the three classes of networks. Dashed red lines correspond to theoretical approximations based on second order motif cumulants, while continuous colored lines correspond to full (exact) values. **e**) Dimensionality versus average correlation.

We find that the dimensionality of the network response for the different connection topologies appears follows the same general trend: it decreases as a function of the average connectivity *p* (cfr. Fig. 5b), until it reaches the boundary of instability for the dynamics. Interestingly, the relation between the average correlation and the dimensionality appears to be very tightly stereotyped as shown in Fig. 5c. Such a tight relationship suggests that one may be able to to interpret average correlations in observed in a circuit in terms of their dimensionality, at least across some classes of network connectivity. Overall, these results suggest that the framework and results given so far do generalize to a more general class of excitatory networks.

In Fig. 6 we move beyond excitatory networks to consider the case of excitatory/inhibitory networks To do so we analyze a random Erdos-Renyi network where 10% of the neurons are randomly selected to be inhibitory and balance out, on average, the excitatory connection weight in the network (see Methods l for more details). The result of this process is a block Erdos-Renyi network with a non-trivial statistics of motifs and cumulants. The sign of the motifs reflects their excitatory, inhibitory or mixed nature. Importantly E-I networks tend to be more stable, which allows for stronger synapses overall. Taking advantage of this we increase the average synaptic strength by changing its scaling from 1*/N* to 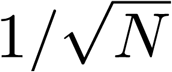 [63, 59].

**Fig 6.**
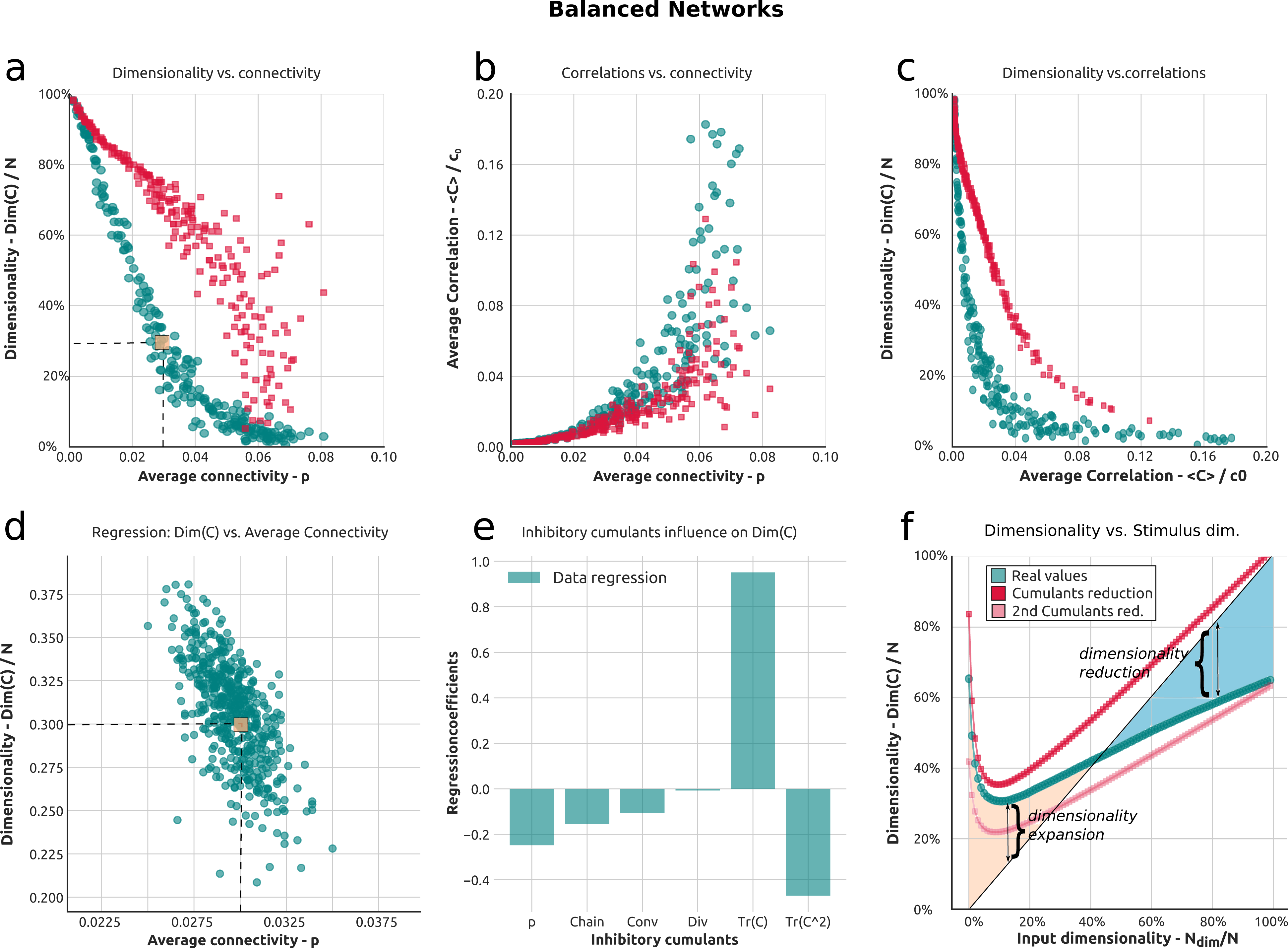
Dimensionality in random balanced recurrent networks and the role of connectivity motifs. **a**) Dimensionality as a function of average connectivity in E-I balanced networks. Highlighted in orange is a network producing low dimensionality. **b**) Average correlation versus average connectivity. **c**) Dimensionality versus average correlation. **e**) Dimensionality as a function of stimulus dimension. The green line corresponds to the full theory, while the red and pink lines correspond to theoretical approximations in the second order cumulant framework, respectively for low and high stimulus dimensionality. The areas in orange and blue indicate, respectively, dimensionality expansion or reduction in the network (cfr. Fig. 4f. **f**) Dimensionality versus average inhibitory connectivity for an ensemble of balanced networks with low dimensionality. The ensemble statistics are averaged around the network corresponding to the orange point. **d**) Regression of the dimensionality against the cumulants of the inhibitory population.

We see that the resulting relationship between the dimensionality of the network and the average synaptic connectivity *p* in Fig. 6a is even stronger that in the fully excitatory case of Fig. 3a. Specifically, Fig. 6a shows that the dimensionality rapidly decreases as a function of the average connectivity, and – different from the purely excitatory case – does so with a very steep initial slope. Moreover, the dimensionality decreases very quickly as a function of average correlations Fig. 6c, so that, once again, E-I networks whose activity might at first appear to be (at least on average) independent due to low values of average pairwise correlations actually show very tightly coordinated dynamics. We also find that the theoretical approximation (red dots), despite capturing the overall steeply decreasing trend, is in poor agreement with the full (exact) values of dimensionality. This is due to the fact that the theory shown is perturbative: we keep terms only up to second order (second order cumulants cfr. Fig. 2). While it could be expanded to account for higher orders at the price of increasing its complexity [18], we here instead highlight its limitations of at second order, at least in our hands, while pointing out important trends in the relationship between the dimensionality of the response and other network properties. For example Fig. 6b and Fig. 6c show that, in the balanced case, the theory approximates to a better extent the average correlations (Fig. 6b) than the dimensionality [17, 18].

To highlight the role of cumulants in controlling these effects we carry out a similar analysis to the one illustrated in Figs. 3d to 3g. We compute the dimensionality for an ensemble of 500 SONETS networks of 1000 neurons each (see Methods m) with excitatory connectivity *p* = 0.03. The average connectivity between inhibitory neurons, together with the motif content, varies perturbatively around *p* = 0.03. How the dimensionality varies as a function of the probability of connection between inhibitory neurons is shown in Fig. 6f, for each network in the ensemble. We then carry out a multilinear regression where the dimensionality of the network is regressed against all the values of the cumulants between neurons in the inhibitory population (*r*^2^=0.420). The result is shown in Fig. 6d. This result is similar to the one shown in Fig. 4e and shows how different motifs may lead to a dimensionality increase or decrease.

One of the main characteristics of E-I balanced networks is the cancellation between strong excitatory and inhibitory contributions. This, in turn, means that the network tends to be in a strongly coupled regime where the internal dynamics is strong and the inputs, rather than driving the network, are entrained to its dynamics. This is shown in Fig. 6e, where the dimensionality of the network varies with the input dimensionality but the span over which the former is modulated by less than 30%, from a dimensionality of roughly 30% to a dimensionality of roughly 60%, over a wide range of input dimensions. If we imagine the input to itself vary in a reasonable range of, say, 30% then the network acts to equalize the dimensionality of its response across this range. Specifically, this seems to be achieved optimally at the minimum of the green line in Fig. 6e, where the contribution of the input and internal network dynamics appears to be of similar strength. This may be an important working point for the network, as we will further cover in the Discussion.

## 3 Discussion

We have introduced a theory of dimensionality in linear, spiking recurrent networks, which predicts the dimensionality of a network’s response from basic features of its internal connectivity and the stimuli it receives. The theory builds on the existing framework of motif cumulants [19, 17, 18, 42], which identified the significance of connectivity motifs in leading a number of other effects in the network dynamics. We single out three important results from our analysis for further discussion here.

First, we find that the statistics of highly local “second order” connectivity motifs – subnetworks of just two or three cells at a time – can be used to predict several (but not all) global aspects of the dimensionality of network-wide activity. These are as follows: for purely excitatory, autonomously spiking networks, the values of connection probability and the prevalance of second order connectivity motifs provides highly accurate quantitative predictions of dimension – and hence dimension appears to be regulated by these connection features alone. For excitatory-inhibitory networks, we can use these localized motifs to make qualitative predictions about trends in dimension with connectivity, but quantitative estimates have large errors. The same is true about the network response to strong inputs: trends can be captured from local motif cumulants, but quantitative accuracy demands a fuller description of network connectivity.

The ability, when it occurs, of local circuit features to regulate global activity patterns is important because local activity dependence appears as one of the major constraints in biological learning paradigms [64, 65, 66]. Thus, when it succeeds, expressing neural dynamics in terms of local connectivity motifs may reveal the function of learning rules, and how they target the dynamics of specific connectivity patterns [67, 68].

Second, our results show that the dimensionality of the network activity has the tendency to assume low values, even when the average pairwise correlations in a network are themselves so low that it might be tempting to consider them as neglibible. In Figs. 3h, 5e and 6c we have shown that, across a number of different connectivity regimes, the network response has low dimensionality when the average correlation is lower than 0.025. This effect is important, it may point to the dimensionality, rather than the often reported statistic of average pairwise correlations (see review in [34]), as a better metric for describing how strongly network activity is coordinated [43, 26]. Moreover, our theory suggests that specific connectivity motifs, i.e. reciprocal motifs, have a prominent role in influencing the activity dimensionality over and above its average correlation.

Third, depending on stimulus properties and network connectivity, the network response may have a higher or lower dimensionality than the stimulus; in this way, feeding a stimulus to a network results in either an expanded or contracted dimensionality in the response (cfr. Figs. 4 and 6e). Which of these occurs depends strongly on the network connectivity. Here, stronger coupling leads to a more restricted range of dimensionalities with which the network operates. This restricted range – produced in response to a wide range of stimuli – may be interpreted as a type of “dimensionality equalization:” the network reduces or expands the stimulus dimensionality to lie in a relatively tight range Figs. 4d and 6e. Moreover, when inputs assume a fixed strength in each dimension, there is a specific stimulus dimensionality where the network response assumes minimum value. This point is of interest as it marks the transition from a dynamical regime dominated by the internal network response to one governed by the stimulus: thus, near the minimum, the network is entrained by the stimulus but not dominated by it, with the internal dynamics serving as scaffold for the activity that is produced.

We close by considering three future research directions that our work here has helped to define.

The first is the question of finding efficient, readily measurable features of network connectivity that drive key aspects of neural network dynamics. Here, we demonstrated some substantial new successes, and failures, of local connectivity motifs in this regard. Further research across our field will be important to understand the relevance of specific connectivity patterns and their statistics, including how they vary across space and cell types [69, 21]. This will be especially interesting in relation to next generation connectomics data, which may unlock new roles and new forms of connectivity structures.

The second is the extension of link between connectivity structure and activity dimension to nonlinear networks. While the theory in this study is for networks that are linearized around their working point, recent work [70, 71] has developed an expansion that predicts that predicts correlations of arbitrary orders in similar Poisson-type networks, for increasing orders of nonlinearity. Further work to elucidate their influence in shaping the dimensionality of neural response would extend the scope of the present analysis beyond linear circuits, possibly bridging our framework with others that have been recently advanced [72].

The third and final direction for future study is analysis of the stimulus entrainment of network dynamics highlighted above. Specifically, neural representations, i.e. the encoding of stimulus-specific information by neural networks, may involve circuitry that either increases, decreases, or equalizes the dimensionality of neural responses, but further work is needed to understand the implications for neural coding [26, 3, 2].

## 4 Methods

### a) Full expression of Dimensionality as a function of cumulants

The expression for the dimensionality *Dim*(***C***) is the ratio between Tr(***C***)^2^ and Tr(***C***^2^). In terms of cumulants the functions of Eq. 11 can be written as:

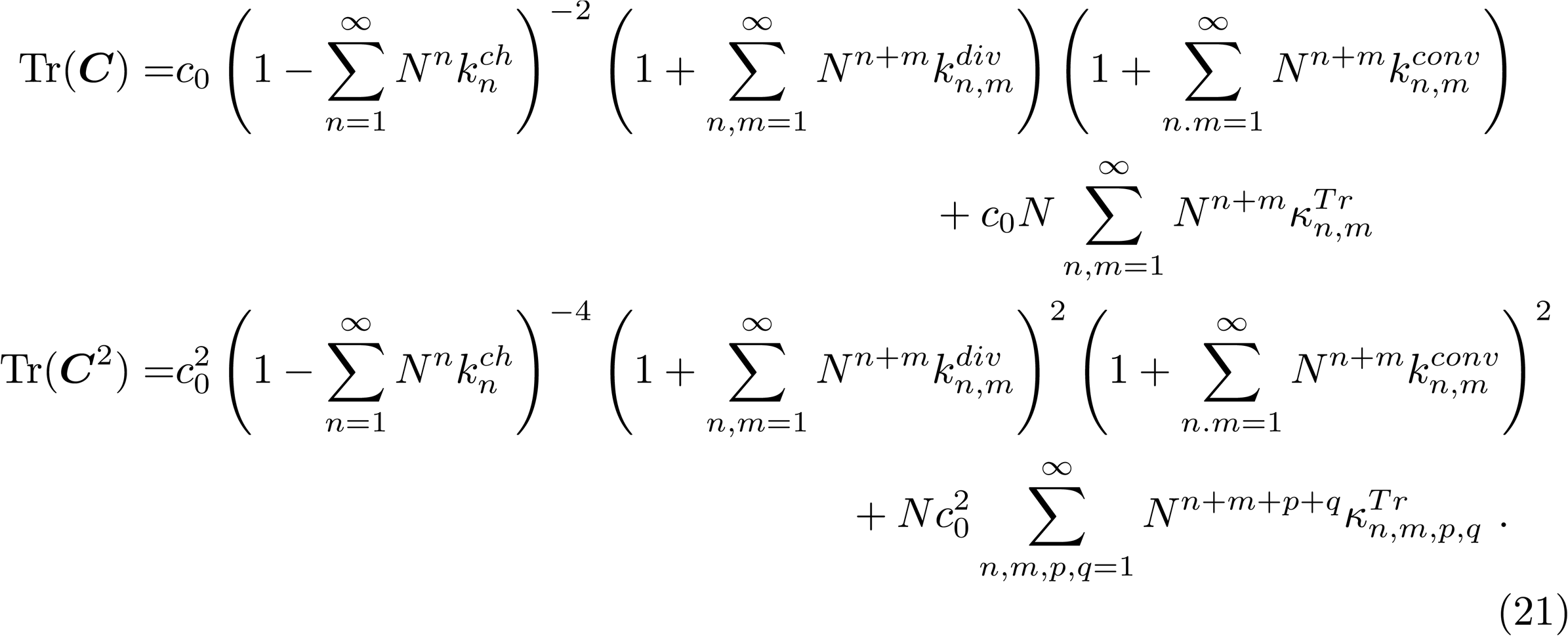

These two expressions are both tightly linked to the average covariance Eq. 10. In particular they can be written as follows to highlight this connection:

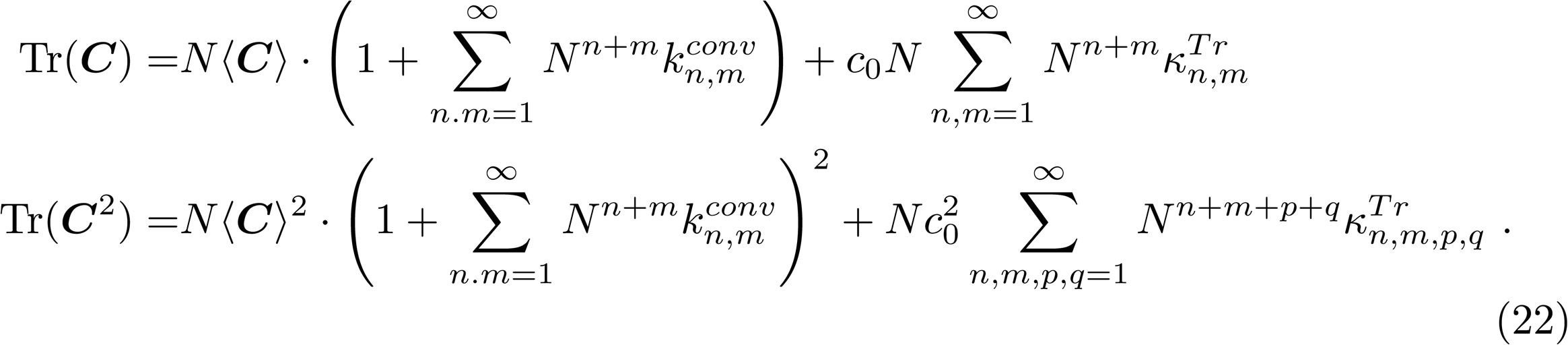

### b) Full expression of dimensionality as a function of cumulants, in the presence of input stimuli

The full expression for Eq. 19 is:

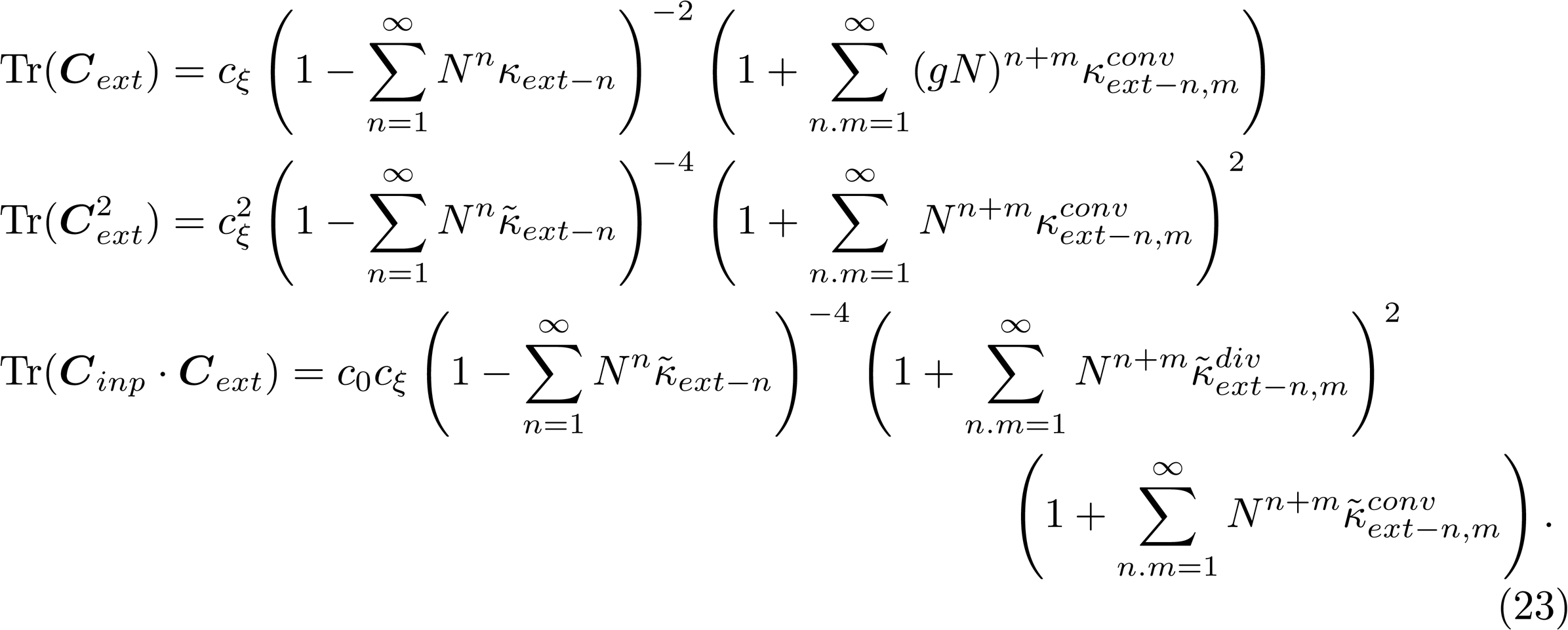

These formulas can be resumed with different choices of cumulants. In particular both ***κint***and ***κext*** can be employed simultaneously (cfr. Suppl.Mat. sec. 3.2). In Eq. 23 we show the expression used to generate figures in the main text; this choice is best motivated in the case of low dimensional input.

### c) Description of SONET networks

The SONET model for random graphs can be seen as an extension of the Erdos-Renyi model. In an Erdos-Renyi graph two nodes are randomly connected with probability *p* (0 ≤ *p* ≤ 1). In SONET networks also second order connection motifs (convergent, divergent, etc.) appear with controlled statistics (see [40, 56] for further details). As an extension of the Erdos-Renyi model, the algorithm we use (provided by the authors of [40]) generates a W with binary entries.

### d) Details for Fig. 1f, Figs. 3b to 3h

The ensemble of networks used for Fig. 3b, Fig. 3c and Fig. 3h consists of 500 networks with *N* = 1000 neurons each. All networks share the parameters *c*_0_ = 1, *A*_*ii*_ = 10 ∀ _*i*_ ∈ {1..*N*} while the connectivity graph *W* is generated through the SONET algorithm with the same set of parameters (*α*’s) used in [40]. Such parameters regulate the statistics of convergent, divergent, reciprocal and chain motifs. They are uniformly sampled in the ranges *p ∈* [0.01, 0.1], *α_recip_ ∈* [-1, 4], *α_conv_* ∈ [0, 1], *α_div_* ∈ [0, 1] and *α_chain_* ∈ [-1, 1].

### e) Details for the regression in Figs. 3d to 3g

The ensemble of networks used for Figs. 3d to 3g has exactly the same parameters as the one above, except that the range for *p* is different: *p* ∈ [0.078, 0.082]. The dimensionality for each network is computed and the difference in dimensionality from an Erdos-Renyi network with *p* = *pER*= 0.08 is regressed against 6 different variables: *p - p_ER_*, and the value of chain, convergent, divergent and the two trace cumulants. The coefficients of the regression are displayed in Fig. 3e.

### f) Details for Figs. 4a, 4c and 4d

These figures display the full covariance dimensionality expression Eq. 3 and the motif reduction Eq. 11 for a SONET network with p=0.03 and a random choice of second order motifs. An input of varying strength and number of factors is fed onto the network. This is captured by

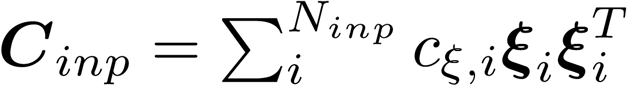 where each ***ξ*** is a random vector of unit norm. In the case of Fig. 4a the number of factors *N*_*dim*_ is increased and *c****_ξ_*** = 0.05. In Fig. 4c the number of factors is one while *c****_ξ_*** is increased. In Fig. 4d the number of factors is increased but the total strength constrained to 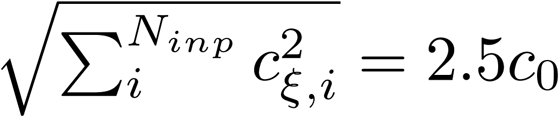.

### g) Details for Figs. 4e, 4g and 4h and theoretical approximation

The procedure for obtaining these figures is equal to the one used for Figs. 4a, 4c and 4d except that the initial network is a SONET network with *p* = 0.08 and random second order motifs.

The pink line in these figures corresponds to a theoretical approximation of the formula in Eq. 18. The term in the denominator Tr(***C*_*int*_· *C*_*ext*_**) is the only term in the expression with the product ***C*_*int*_** and ***C*_*ext*_**. We used the following inequality to build an upper bound for this term:

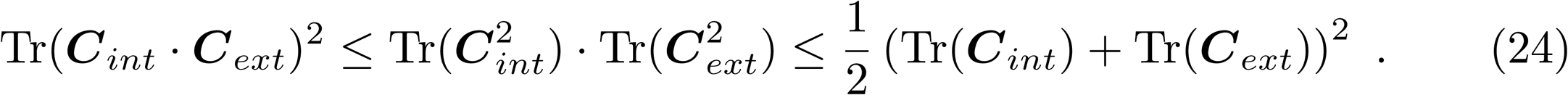

By substituting the rightmost side of this expression into Eq. 18 we obtain the expression for the pink line displayed in Figs. 4a, 4c and 4d.

### h) Details for Figs. 5a to 5e

The figures use the same values and techniques of Figs. 3b and 3c. The different network architectures are all generated using the package NetworkX in Python 3.6. The Erdos-Renyi network is a randomly connected network, the small world network has a number of nodes denoted by *p · N* and probability of rewiring 0.3, the scale-free network is obtained through a Barabasi-Albert graph where the number of number of edges to attach from a new node to existing nodes (parameter *m*) is derived as a function of the final number of connections *p · N* and the number of nodes 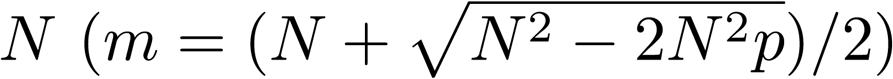.

### i) Details for Figs. 6a to 6e

The networks displayed in these figures are 500 SONET networks with average synaptic strength 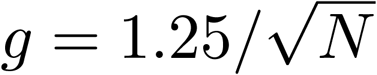 that scales with 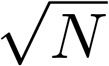 rather than *N*. For each network a random 10% of the neurons is selected to be inhibitory and their strength rescaled so that ⟨***G*_*EE*_**⟩ = ⟨***G*_*II*_**⟩ where ***G*_*EE*_ and *G*_*II*_** denote respectively the part of the connectivity graph ***G*** in between the excitatory and the inhibitory population. We checked that the network so obtained respects the constraints for a balanced state determined in [73].

### j) Details for Figs. 6d and 6f

We generate 500 SONET networks with connectivity *p* = 0.03. Upon balancing the network 10% of the neurons are inhibitory. The dimensionality of this ensemble of networks is regressed against the values of the connectivity cumulants computed on the inhibitory part of the network.

## Acknowledgments

This work was supported in part by NSF grant DMS-1514743. We are grateful to Yu Hu and Kameron Harris for helpful insights and comments. We thank the Allen Institute founders, Paul G. Allen and Jody Allen, for their vision, encouragement and support.

